# Glioblastoma Hijacks Astrocyte Ribosomes to Feul Malignancy through Translatome Reprogramming

**DOI:** 10.64898/2026.10.28.762373

**Authors:** Xujia Wu, Chen Zheng, Lanjie Chen, Junju Chen, Yuchao Zhang, Long Cheng, Yushan Peng, Xuechao Zeng, Mingpu Gao, Feng Luo, Yanxing Lai, Jiying Cheng, Yixin Gao, Jian Zhong, Xuesong Yang, Nunu Huang, Feizhe Xiao, Xuesong Liu, Fan Bai, Nu Zhang

## Abstract

Cancer cells demand elevated protein synthesis, yet whether they directly requisition ribosomes from microenvironmental host cells remains unknown. Here, utilizing *in viv*o and *in vitro* ribosome tracing, we show that glioblastoma (GBM) cells hijack ribosomes from surrounding astrocytes via tunneling nanotube-like structures. This transfer supports GBM malignancy in both murine and patient-derived models. Single-cell RNA sequencing along with whole-cell and cell-source-traceable ribosome profiling reveal that transferred ribosomes are translationally active and selectively reprogram the tumor translatome, prioritizing cell-cycle genes and driving interferon-response-mediated immune evasion. Through a genome-wide CRISPR screen, we identify INTS12 as a critical mediator of ribosome transfer. Genetic inactivation or AAV-mediated knockdown of INTS12 impairs ribosome uptake, curbs translation and proliferation, remodels the immunosuppressive landscape, and extends survival in orthotopic models. Our study uncovers a parasitic translation-support mechanism in GBM, establishing INTS12 as a therapeutic vulnerability to disrupt ribosome transfer.

## INTRODUCTION

Cancer cells co-opt surrounding non-malignant cells-‘‘accessories to the crime”-to support their own growth^1^. One common strategy involves intercellular communication and material transfer, which includes DNA, RNA, proteins, small metabolites, or even entire organelles such as mitochondria and lysosomes, occurring either indirectly via extracellular vesicles^2–4^ or directly through intercellular conduits like tunneling nanotubes (TNTs) or tumor microtubes (TMs)^5–12^. Tumor proliferation and progression demand high rates of protein synthesis, which are often sustained by cell-autonomous activation of the translation machinery and enhanced ribosome biogenesis^13,14^. However, whether cancer cells can directly hijack ribosomes from neighboring normal cells to meet their translational needs remains elusive.

Glioblastoma (GBM) is a lethal brain cancer, featuring rapid proliferation and pronounced immunosuppression, and carries a median survival of less than two years^15,16^. Within the brain tumor microenvironment, astrocytes are the most abundant stromal cells. They normally preserve homeostasis and support neighboring cells under physiological conditions, but often proliferate and undergo reprogramming into reactive astrocytes in response to pathological injury and inflammation, thereby actively shaping neurological disease progression^17–19^. Accumulating evidence indicates that they support malignant proliferation and immunosuppression through diverse mechanisms, including cytokine secretion and direct cell-cell interactions with tumor cells and immune cells^20–24^. Astrocytes have been shown to transfer functional mitochondria to tumor cells^8,12^. However, whether they also donate ribosomes has yet to be explored.

In this study, we observed widespread ribosome transfer from astrocytes to GBM cells via TNT-like structures. Transferred ribosomes were functional and actively engaged in protein translation, reprogramming the tumor translatome toward proliferation and interferon (IFN) response, thereby accelerating tumor growth and mimicking a sustained IFN-stimulated state that fosters an immunosuppressive microenvironment. A genome-wide CRISPR knockout screen identified INTS12 as a critical mediator of ribosome transfer. Targeting INTS12 suppressed ribosome uptake, reversed translation reprogramming, restrained GBM proliferation, and remodeled the immune microenvironment into an antitumor state, offering a potential therapeutic paradigm for this devastating disease.

## RESULTS

### Glioblastoma hijacks ribosomes from peripheral brain

To sustain their voracious energy requirements, GBM cells acquire functional mitochondria from the microenvironment^8,11,12^. Considering the parallel demand for protein synthesis in this lethal cancer, we reasoned that ribosomes might be acquired through a similar intercellular mechanism. To test this hypothesis, we generated a conditional reporter mouse model expressing a green fluorescent protein (GFP) tagged ribosomal protein RPL10A, designated "RiboTracer" (Loxp-stop-loxp [LSL]-Avi-GFP-*Rpl10a*) (Figure S1A). In this system, Cre-mediated recombination induces expression of the GFP-RPL10A fusion protein, which integrates into intact ribosomes and label translationally active ribosomes with green fluorescence^25,26^. To achieve ubiquitous ribosome labeling, we crossed the RiboTracer allele with *UBC*-Cre-ERT2^27^, where a broadly active promoter directs tamoxifen-inducible Cre-mediated recombination, resulting in pervasive labeling (Figure S1B). We then orthotopically implanted mKate2 labeled brain tumor cells (GL261 or CT2A) into these mice. In the resulting tumors, we observed widespread green fluorescence within the red-labeled tumor cells (Figures 1A and 1B). By contrast, in control mice carrying the RiboTracer allele without the *UBC*-Cre-ERT2 driver, no GFP signal was detected in either tumor cells or surrounding brain tissue, confirming that the observed green fluorescence was strictly dependent on Cre-mediated recombination and did not arise from leaky reporter expression or non-specific fluorescence (Figures 1A and 1B). Flow cytometry analysis revealed that many GBM cells contained detectable labeled ribosomes (Figures 1C and 1D; gating strategy as in S1C). These findings suggest that GBM cells broadly acquire ribosomes from surrounding normal brain *in vivo*.

**Figure 1.**
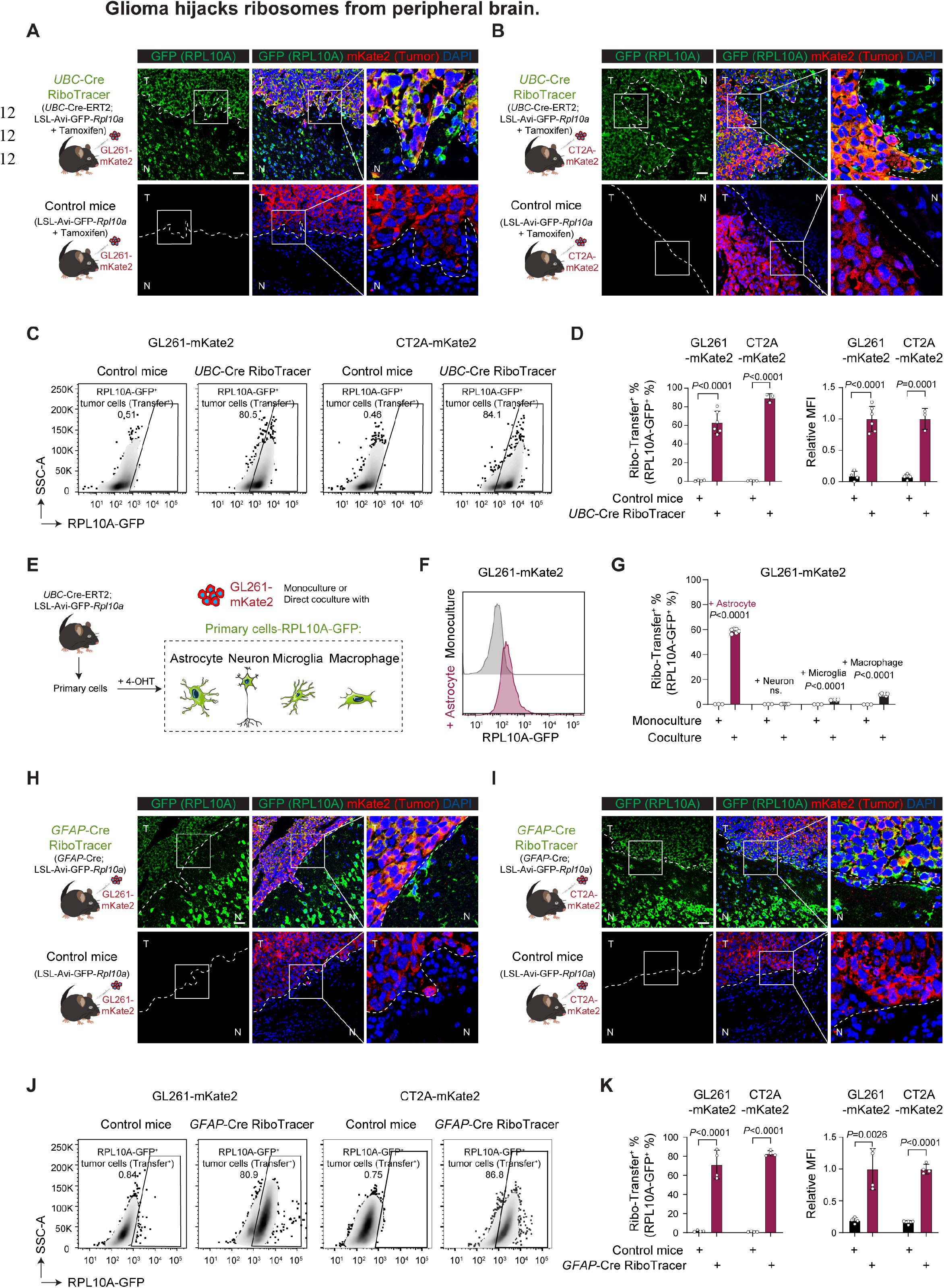
Glioma hijacks ribosomes from peripheral brain. (**A** and **B**) Schematics and representative images of RPL10A-GFP in brain slices from GL261- or CT2A-bearing *UBC*-Cre; RiboTracer mice and controls. mKate2 labels tumor. T, tumor. N, normal. Scale bar, 50 μm. (**C** and **D**) Representative flow plots and quantification of of RPL10A-GFP in GL261 and CT2A GBM cells isolated from orthotopic tumors of *UBC*-Cre; RiboTracer mice and controls (n=4 for each control group, n=6 for GL261 and n=3 for CT2A in *UBC*-Cre; RiboTracer group). (**E-G**) Schematics, representative flow plots, and quantification of Ribo-Transfer percentage in GL261 cells monocultured or directly cocultured with indicated cells (n=3 independent experiments for monoculture, n=6 for coculture). (**H** and **I**) Schematics and representative images of RPL10A-GFP in brain slices from GL261- or CT2A-bearing *GFAP*-Cre; RiboTracer mice and controls. mKate2 labels tumor. T, tumor. N, normal brain. Scale bar, 50 μm. (**J** and **K**) Representative flow plots and quantification of of RPL10A-GFP in GL261 and CT2A GBM cells isolated from orthotopic tumors of *GFAP*-Cre; RiboTracer mice and controls (n=4 per group). Data are represented as the mean ± SD in (D), (G) and (K). Individual data points represent one mouse unless otherwise specified. Two-tailed unpaired t test for (D), (G) and (K). ns, not significant. See also Figure S1.

The cellular landscape of the brain tumor microenvironment comprises multiple non-malignant lineages^28^. To determine the primary donor of ribosomes to GBM cells, we isolated major tumor-infiltrating cell types from reporter mice and directly cocultured these cells with mKate2^+^ GBM cells (Figure 1E). Among these cells, astrocytes exhibited the highest capacity for ribosome transfer, whereas myeloid cells contributed only minimally and neurons showed no detectable transfer (Figures 1F and 1G), supporting astrocytes as the major ribosomal donors to tumor cells. To examine whether astrocytic ribosome transfer occurs *in vivo*, we crossed the RiboTracer allele with *GFAP*-Cre, which is commonly used to label astrocytes^29–31^. Immunofluorescence staining of brain sections from tumor-free mice confirmed that GFP-RPL10A signals were specifically colocalized with GFAP positive astrocytes, whereas no such signals were observed in control mice lacking the *GFAP*-Cre driver (Figure S1D), validating the specificity of this labeling system. Following orthotopic implantation of mKate2⁺ GBM cells, we detected astrocyte-derived ribosome fluorescence in the majority of tumor cells (Figures 1H-1K). In control mice lacking the *GFAP*-Cre driver, no such green fluorescence was observed in tumor cells (Figures 1H and 1I), confirming that the signal originated specifically from Cre-mediated recombination and subsequent ribosome labeling in astrocytes. Together, these results establish that GBM cells hijack ribosomes from peripheral cells, with astrocytes serving as the primary donor.

### Ribosomes are directly transferred through tunneling nanotube-like structures

To test whether ribosome transfer also occurs in patient-derived tumor cells, we established GFP-labeled patient-derived GBM cells and directly co-cultured them with a human astrocyte expressing RPL10A-mKate2. Astrocytic RPL10A signals were readily detected within the GBM cells (Figures 2A-2C; gating strategy as in Figure S2A). To rule out false positives from a single tag, we independently labeled astrocytic ribosome with RPL36-mKate2, a distinct ribosomal protein that incorporates into translating ribosomes^32^. Comparable transferred ribosome signals were detected in tumor cells (Figures 2D and S2B). To exclude the possibility that the observed signals merely reflect nonspecific transfer of fluorescent proteins rather than bona fide ribosome transfer, we labeled tumor cells with GFP and astrocytes with mKate2 alone. In direct co-culture experiments, no significant fluorescence transfer was detected (Figures S2C-S2E). These results demonstrate intercellular ribosome transfer in patient-derived GBM cells.

**Figure 2.**
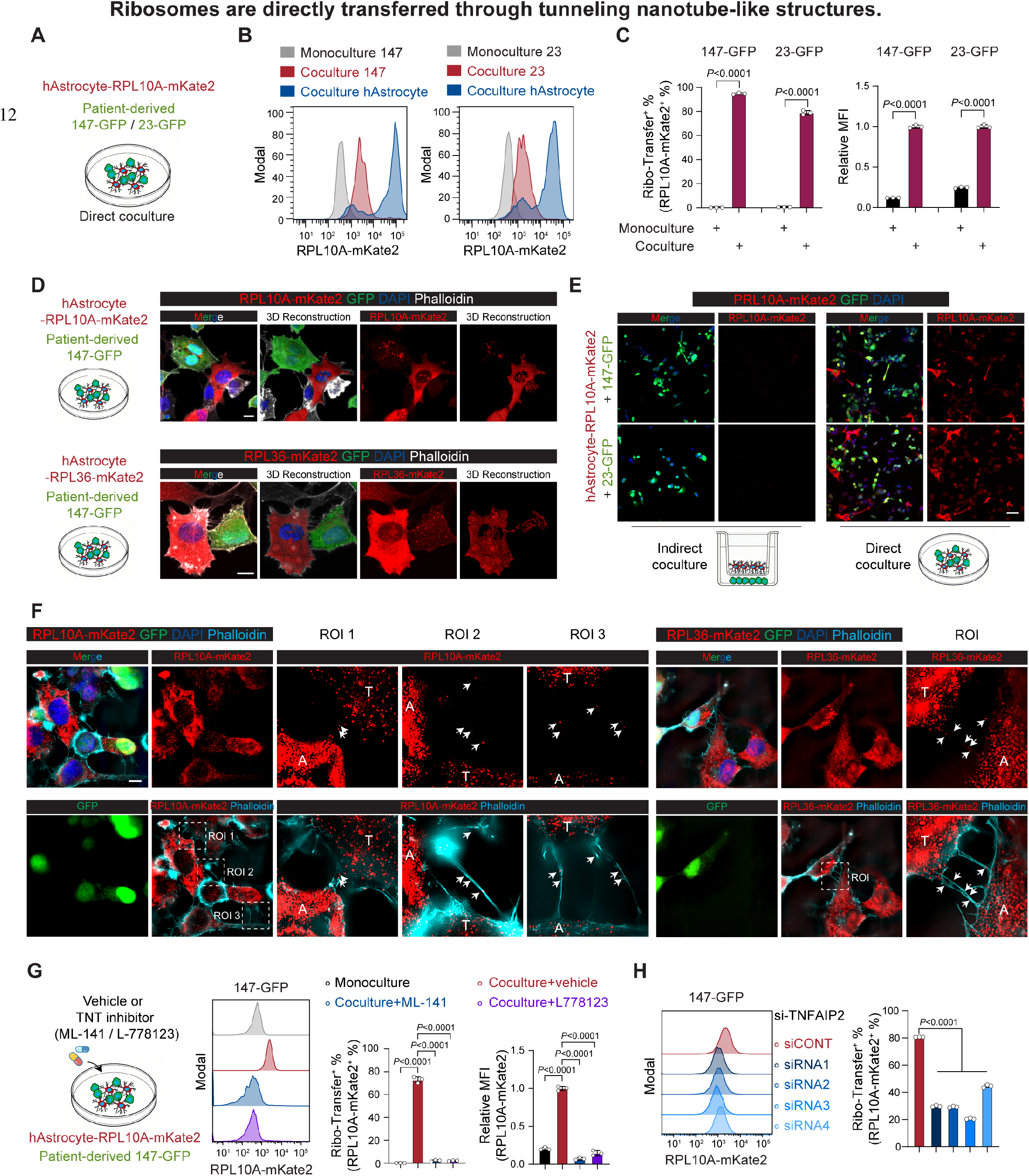
Ribosomes are directly transferred through tunneling nanotube-like structures. (**A-C**) Schematics, representative flow plots, and quantification of RPL10A-mKate2 in GFP- labeled patient-derived #23 or #147 GBM cells cultured alone or directly cocultured with human astrocytes expressing RPL10A-mKate2 (n=3 per group). (D) Schematics and representative images of RPL10A-mKate2 or RPL36-mKate2 in the indicated coculture system. GFP labels tumor cells; phalloidin labels F-actin (TNT marker). Scale bar, 10 μm. (E) Schematics and representative images of RPL10A-mKate2 in indirect coculture separated by a transwell chamber or in direct coculture. Scale bar, 50 μm. (F) Representative super-resolution confocal images of RPL10A-mKate2 or RPL36-mKate2 in direct coculture. ROI, enlarged region of interest. GFP labels tumor cells; phalloidin labels F-actin (TNT marker). A, astrocyte. T, tumor cell. White arrows indicate ribosome particles within TNT- like structures. Scale bar, 10 μm. (G) Schematics, representative flow plots, and quantification of RPL10A-mKate2 in direct coculture treated with 10 μM ML-141, 10 μM L-778123, or vehicle (n=3 per group). (H) Representative flow plots and quantification of RPL10A-mKate2 in direct coculture of human astrocyte expressing RPL10A-mKate2 and 147-GFP treated with the indicated siRNAs targeting *TNFAIP2* (n=3 per group). Data are represented as the mean ± SD in (C), (G) and (H). Individual data points represent independent experiments. Two-tailed unpaired t test for (C). One-way ANOVA followed by multiple comparisons for (G) and (H). See also Figures S2 and S3.

Intercellular organelle transfer can occur either indirectly via extracellular vesicles or directly through intercellular conduits like TNTs or TMs^7,8,11,12^. To determine the mechanism underlying astrocytic ribosome transfer, we co-cultured astrocytes and tumor cells either in direct contact or separated by a transwell chamber. Ribosome transfer was detected only under direct co-culture conditions (Figures 2E, S2F and S2G), indicating a requirement for cell-cell contact during transfer. Supporting this, super-resolution confocal microscopy revealed transferring ribosome particles within TNT-like structures (Figure 2F), identified by well-established phalloidin staining of F-actin^7^. Rho GTPases are central regulators of actin reorganization and their pharmacological inhibitors ML-141 (targets Cdc42), and L-778123 (targets farnesyltransferase and geranylgeranyltransferase type 1) has been shown to affect the assembly of TNT-like structures^6^. Treatment with these inhibitors reduced ribosome uptake by GBM cells (Figures 2G and S2H). TNFAIP2 (also known as M-sec) is a key driver of TNT-like structures^33^. Knockdown of TNFAIP2 in GBM cells reduced astrocytic ribosome transfer (Figures 2H, S2I and S2J). These results establish that TNT-like structures mediate ribosome transfer from astrocytes to GBM cells.

TNT-like structures serve as bidirectional conduits, allowing passive diffusion or directed uptake of cargo^7^. To determine the direction of ribosome transfer, we performed reciprocal co-culture experiments by labeling ribosomes in tumor cells instead of astrocytes (Figure S3A). Under these conditions, we did not observe significant ribosome uptake by astrocytes (Figures S3B and S3C). To test this *in vivo*, we implanted RPL10A-GFP-expressing GBM cells into the brains of tdTomato reporter mice. Ribosome signals derived from tumor cells were rarely detected within tdTomato⁺ non-malignant cells (Figure S3D). These results indicate that ribosome transfer is predominantly unidirectional, with tumor cells actively acquiring ribosomes from surrounding astrocytes, suggesting a potential addiction of GBM cells to this exogenous ribosome supply.

### Ribosome transfer fuels cancer malignancy

Given the pronounced ribosome hijacking from astrocytes by GBM cells, we hypothesized that this ribosome transfer might support tumor malignancy. To test this, we orthotopically implanted mKate2-labeled GBM cells into *GFAP*-Cre; RiboTracer mice. At the endpoint, tumor cells were sorted into Ribo-Transfer^lo^ and Ribo-Transfer^hi^ populations based on the transferred RPL10A-GFP intensity (Figures 3A and 3B). Global protein synthesis rates were measured using *O-*propargyl-puromycin (OPP), a puromycin analog widely used to monitor translational activity^34,35^. Ribo- Transfer^hi^ GBM cells exhibited higher OPP incorporation than Ribo-Transfer^lo^ cells (Figures 3C and 3D), indicating activation of global translation. These two populations were isolated and used for secondary tumor implantation. Ribo-Transfer^hi^ cells gave rise to more aggressive tumors, characterized by accelerated growth and reduced host survival (Figures 3E-3G). The enhanced translational phenotype was maintained, as secondary tumors derived from Ribo-Transfer^hi^ cells showed stronger OPP incorporation (Figure 3H). In direct co-cultures of patient-derived GBM cells and human astrocytes, Ribo-Transfer^lo^ and Ribo-Transfer^hi^ GBM populations were isolated in a similar way (Figures 3I and 3J). Consistently, Ribo-Transfer^hi^ population exhibited more active protein synthesis, as evidenced by increased OPP incorporation and puromycin labeling (Figures 3K and 3L). Secondary transplantation of these patient-derived populations into immunocompromised mice revealed that Ribo-Transferʰⁱ cells formed more devastating tumors, with accelerated growth (Figures 3M and 3N), shortened host survival (Figure 3O), and sustained high translational activity (Figures 3P and 3Q), recapitulating the malignant phenotype observed in the mouse model. These results indicate that astrocytic ribosome transfer robustly enhances protein synthesis and fuels malignant progression in GBM.

**Figure 3.**
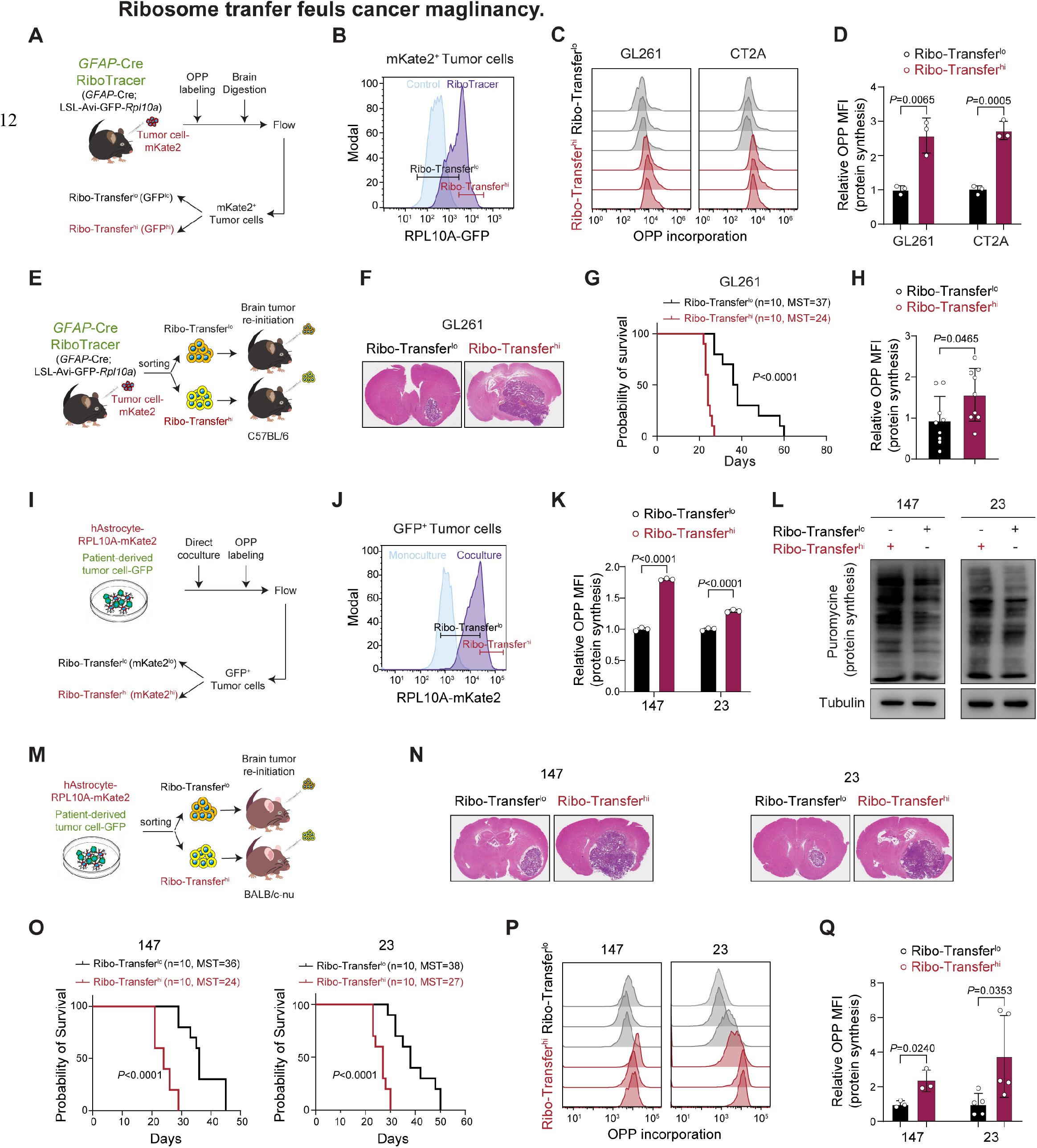
Ribosome transfer fuels cancer malignancy. (**A** and **B**) Schematics and representative flow plots of the isolation strategy for Ribo-Transfer^lo^ and Ribo-Transfer^hi^ murine tumor cells in *GFAP*-Cre; RiboTracer mice. (**C** and **D**) Representative flow plots and quantification of OPP incorporation in Ribo-Transfer^lo^ and Ribo-Transfer^hi^ GL261 or CT2A cells (n=3 per group). (**E**) Schematics of brain tumor re-initiation by Ribo-Transfer^lo^ and Ribo-Transfer^hi^ tumor cells isolated from *GFAP*-Cre; RiboTracer mice. (**F** and **G**) Representative HE-stained brain sections (F, collected at the onset of neurological symptoms in any group) and Kaplan–Meier survival curves of tumor-bearing mice (G, n=10 per group) implanted with Ribo-Transfer^lo^ or Ribo-Transfer^hi^ GL261 cells. MST, median survival time. (**H**) Quantification of OPP incorporation in brain tumors initiated by Ribo-Transfer^lo^ and Ribo- Transfer^hi^ GL261 cells (n=9 per group). (**I** and **J**) Schematics and representative flow plots of the isolation strategy for Ribo-Transfer^lo^ and Ribo-Transfer^hi^ patient-derived tumor cells in a direct coculture system. (**K**) Quantification of OPP incorporation in Ribo-Transfer^lo^ and Ribo-Transfer^hi^ patient-derived #23 or #147 GBM cells (n=3 independent experiments per group). (**L**) Immunoblots showing puromycin incorporation of Ribo-Transfer^lo^ and Ribo-Transfer^hi^ #23 or #147 GBM cells. (**M**) Schematics of brain tumor re-initiation by Ribo-Transfer^lo^ and Ribo-Transfer^hi^ tumor cells isolated from the direct coculture system. (**N** and **O**) Representative HE-stained brain sections (N, collected at the onset of neurological symptoms in any group) and Kaplan–Meier survival curves of tumor-bearing mice (O, n=10 per group) implanted with Ribo-Transfer^lo^ or Ribo-Transfer^hi^ #23 or #147 GBM cells. MST, median survival time. (**P** and **Q**) Representative flow plots and quantification of OPP incorporation in brain tumors initiated by Ribo-Transfer^lo^ and Ribo-Transfer^hi^ #23 or #147 GBM cells (n=3 for #147; n=5 for #23). Data are represented as the mean ± SD in (D), (H), (K) and (Q). Individual data points represent one mouse unless otherwise specified. Two-tailed unpaired t test for (D), (H), (K) and (Q). Log-rank test for (G) and (O).

### Ribosome transfer rewires cancer translation to drive proliferation and immune evasion

To elucidate the mechanisms by which ribosome transfer supports GBM malignancy, we sorted Ribo-Transfer^lo^ and Ribo-Transfer^hi^ tumor cells from three tumor-bearing *GFAP*-Cre; RiboTracer mice and performed single-cell RNA sequencing (scRNA-seq) (Figure 4A). After quality control, we obtained a total of 52,556 high-quality cells, which were resolved into five glioma clusters (clusters 0, 1, 2, 3, and 5), along with minor populations of fibroblasts, endothelial cells, and tumor-associated myeloid cells (Figures 4B and S4A). Cluster 0 glioma exhibited high expression of mitochondrial respiratory chain genes (e.g., *Sdhaf1*) and enriched oxidative phosphorylation. Cluster 1 glioma showed features of central nervous system development and angiogenesis. Cluster 2 glioma was proliferative, expressing *CDK1* and *Mki67*. Cluster 3 glioma was associated with inflammatory response and displayed a strong IFN response signature. Cluster 5 glioma was enriched in hypoxia and glycolysis pathways, alongside robust mTORC1 signaling and cholesterol metabolism (Figures S4B and S4C).

**Figure 4.**
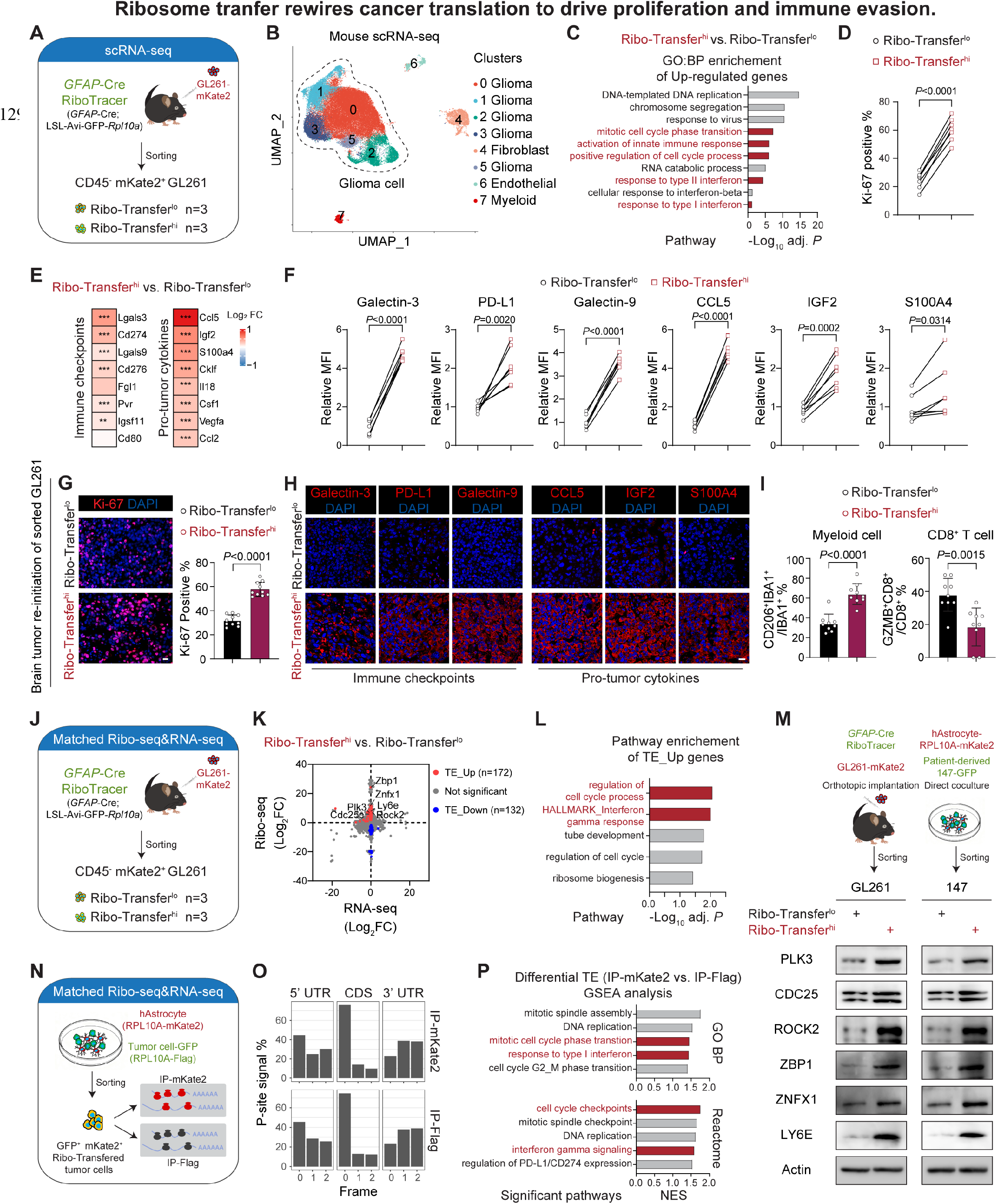
Ribosome transfer rewires cancer translation to drive proliferation and immune evasion. (A) Schematics of scRNA-seq of Ribo-Transfer^lo^ and Ribo-Transfer^hi^ GL261 cells isolated from tumor-bearing *GFAP*-Cre; RiboTracer mice. (B) UMAP plot showing the identified cell clusters from (A). (C) GO:BP enrichment of upregulated genes in Ribo-Transfer^hi^ vs. Ribo-Transfer^lo^ glioma cells in scRNA-seq. (D) Quantification of Ki-67^+^ percentage in Ribo-Transfer^lo^ and Ribo-Transfer^hi^ GL261 cells by flow cytometry (n=7 per group). (E) Heatmap of log2 fold change for indicated genes in Ribo-Transfer^hi^ vs. Ribo-Transfer^lo^ glioma cells from scRNA-seq. (F) Quantification of indicated immune checkpoints and pro-tumor cytokines in Ribo-Transfer^lo^ and Ribo-Transfer^hi^ GL261 cells by flow cytometry (n=7 per group). (**G-I**) Representative staining and quantification of Ki-67^+^ percentage (G, n=10 randomly selected fields from at least three mice), representative staining of indicated immune checkpoints and pro- tumor cytokines (H), and quantification of suppressive myeloid cells (IBA1^+^CD206^+^) and effector T cells (CD8^+^GZMB^+^) (I, n=9 randomly selected fields from at least three mice) in brain sections from tumor-bearing mice implanted with sorted Ribo-Transfer^lo^ or Ribo-Transfer^hi^ GL261 cells. Scale bar, 20 μm. (J) Schematics of matched Ribo-seq and RNA-seq of Ribo-Transfer^lo^ and Ribo-Transfer^hi^ GL261 cells isolated from tumor-bearing *GFAP*-Cre; RiboTracer mice. (K) Four-quadrant plot showing log2 fold change of Ribo-seq and RNA-seq from (J). TE, translation efficiency. (L) GO:BP and Hallmark pathway enrichments of genes with upregulated TE in Ribo-Transfer^hi^ vs. Ribo-Transfer^lo^ GL261 cells. (M) Schematics and immunoblots showing indicated proteins in Ribo-Transfer^lo^ and Ribo- Transfer^hi^ GL261 cells isolated from orthotopic tumors and #147 cells isolated from coculture. (N) Schematics of matched Ribo-seq and RNA-seq using a cell-source-traceable dual labeling system. IP-mKate2 enriches astrocytic ribosomes; IP-Flag enriches tumor ribosomes. (O) Percentage of P-site signals on three-nucleotide (0, 1, 2) codon frames in 5’UTR, CDS and 3’UTR in data from (N). (P) GSEA analysis of GO:BP and Reactome pathways in differential TE analysis of IP-mKate2 vs. IP-Flag. Data are represented as the mean ± SD in (G) and (I). Individual data points represent one mouse unless otherwise specified. Overrepresentation test corrected by FDR for (C) and (L). Two-tailed paired t test for (D) and (F). Two-tailed unpaired t test for (G) and (I). See also Figure S4-S6.

To characterize ribosome transfer in GBM cells, we first compared the cluster composition of Ribo-Transferˡᵒ and Ribo-Transferʰⁱ tumor cells. Although Ribo-Transferʰⁱ cells showed a slight increase in cluster 3 and a slight decrease in cluster 0, both populations were broadly distributed across all clusters (Figure S4D), indicating that the Ribo-Transferʰⁱ phenotype is not restricted to any particular cluster. Differential expression and gene ontology (GO) enrichment analyses revealed the most prominently upregulated pathways in Ribo-Transferʰⁱ GBM cells were cell cycle progression and IFN response (Figures 4C and S4E), reflecting a concurrent activation of proliferation and inflammatory signatures. In contrast, downregulated genes were associated with central nervous system development (Figure S4F). Gene set scoring indicated that Ribo-Transferʰⁱ cells exhibited more active proliferation and IFN response signatures (Figure S4G). In line with the sequencing data, we observed a higher proportion of Ki-67 positive cells among Ribo-Transferʰⁱ population in tumor (Figure 4D). Patient-derived Ribo-Transferʰⁱ GBM cells consistently showed faster proliferation (Figure S5A) and increased 5-ethynyl-2′-deoxyuridine (EdU) incorporation (Figures S5B and S5C). Persistent IFN and inflammatory signaling in tumors trigger both immune checkpoint upregulation and pro-tumor cytokine secretion, collectively reinforcing myeloid suppression and T cell dysfunction, ultimately nurturing an immunosuppressive milieu^36–38^. We found that Ribo-Transferʰⁱ cells upregulated multiple immune checkpoints, including Galectin-3 (encoded by *Lgals3*), PD-L1 (encoded by *CD274*), and Galectin-9 (encoded by *Lgals9*), as well as pro-tumor cytokines such as CCL5, IGF2, and S100A4 (Figures 4E and 4F). In secondary tumors derived from Ribo-Transferʰⁱ cells, both the proliferative and immunoevasive phenotypes persisted (Figures 4G, 4H and S5D). These tumors exhibited an increased abundance of IBA1^+^CD206^+^ immunosuppressive myeloid cells and a reduced presence of CD8^+^GZMB^+^ effector T cells (Figures 4I, S5E and S5F), indicating a tumor-promoting, immunosuppressive microenvironment.

To exclude the possibility that the observed upregulation of proliferation and IFN response signatures was driven by a specific cluster, we profiled these signatures within each glioma cluster. Proliferation signature was consistently upregulated in Ribo-Transferʰⁱ cells across all clusters (Figure S5G). IFN response signatures were primarily upregulated in clusters 0, 1, and 3, which together accounted for approximately 85% of tumor cells (Figures S5H and S5I). These findings suggest that the pathways enriched in Ribo-Transferʰⁱ GBM cells reflect a general alteration rather than a cluster-specific contribution. GBM exhibits pronounced cellular heterogeneity and plasticity, commonly classified into four transcriptional states: OPC-like, NPC-like, AC-like and MES-like^39^. Both Ribo-Transferʰⁱ and Ribo-Transferˡᵒ cells were distributed across all four states with no apparent difference in their distribution patterns (Figure S5J), suggesting that the observed pathway activations are not driven by a preferential enrichment in any particular state.

To determine whether the transferred ribosomes directly drive the phenotypic shift of Ribo-Transferʰⁱ cells, we isolated different transfer populations from tumors and performed ribosome profiling (Ribo-seq), a powerful tool in translatome analysis^40^ (Figure 4J). Translational efficiency (TE) analysis revealed that Ribo-Transferʰⁱ cells exhibited increased translation rates of transcripts enriched in cell cycle regulation and IFN response (Figures 4K and 4L), whereas downregulated TE was primarily associated with neurodevelopment (Figure S6A), suggesting that the Ribo-Transferʰⁱ phenotype may be driven by alterations in the translatome. Consistently, both murine and patient-derived GBM cells showed upregulated translation of core genes involved in proliferation (e.g., PLK3, CDC25, ROCK2)^41,42^ and IFN response (e.g., ZBP1, ZNFX1, LY6E)^43–45^, with increased protein levels but stable RNA levels (Figures 4M, S6B and S6C). These findings suggest that core translational events propel the global phenotypic reprogramming that underpins the aggressive and immunosuppressive traits of Ribo-Transferʰⁱ GBM cells.

To gain deeper insight into the contribution of astrocyte-derived ribosomes to translatome rewiring in Ribo-Transferʰⁱ cells, we developed cell-source-traceable ribosome profiling using a dual-labeling system that resolves the origin of ribosomes within transferred cells (Figures 4N and S6D). In this system, astrocytes expressed RPL10A-mKate2 to mark astrocytic ribosomes, and tumor cells expressed RPL10A-Flag to label their own ribosomes. Following direct co-culture, we isolated ribosome transferred tumor cells and performed immunoprecipitation (IP)-coupled Ribo-seq together with matched RNA-seq. Specifically, IP-mKate2 captured astrocyte-derived ribosomes, whereas IP-Flag captured tumor-derived ribosomes. Astrocyte-derived ribosomes exhibited a strong three-nucleotide periodicity in CDS region similar to tumor-derived ribosomes (Figures 4O and S6E), which is a characteristic of translating ribosomes in Ribo-seq data^40^ and indicates that transferred ribosomes are actively engaged in mRNA translation. Comparison of the translatomes of exogenous and endogenous ribosomes revealed that astrocyte-derived ribosomes were preferentially engaged in translating genes involved in proliferation and the IFN response (Figures 4P and S6F). Astrocytes transition to a reactive state in response to pathological insult. In the context of brain tumors that mimic a chronic form of brain injury, tumor-associated astrocytes within this niche frequently activate proliferative programs and display robust IFN response signatures^46–49^. Together with these findings, our observations support a model in which astrocyte-donated ribosomes not only actively participate in translation but also convey their intrinsic translational preferences to tumor cells, thereby promoting tumor cell proliferation and IFN response, and ultimately contributing to the formation of a pro-proliferative and immunosuppressive tumor microenvironment.

### INTS12 is a targetable regulator of ribosome transfer

Given the potential role of ribosome transfer in driving GBM proliferation and immunosuppression, we sought to examine whether a similar phenomenon exists in clinical samples. We collected tumor samples from 20 patients with IDH-wildtype GBM and performed scRNA-seq (Figure 5A). After quality control, we obtained 189,319 high-quality cells, including 75,559 tumor cells, as well as oligodendrocytes, myeloid cells, neutrophils, B cells, and T cells (Figures 5B and S7A-S7C). Based on transcriptional similarity to mouse tumor cells with defined ribosome transfer status, we predicted 37,779 Ribo-Transferˡᵒ and 37,780 Ribo-Transferʰⁱ GBM cells within the patient samples (Figure S7D). Consistently, patient-derived Ribo-Transferʰⁱ cells exhibited higher scores for translation (Figure S7E), proliferation and IFN response signatures (Figure 5C). We then ranked the 20 patients by the proportion of Ribo-Transferʰⁱ cells and stratified them into low-transfer (Patient_lo) and high-transfer (Patient_hi) groups (Figure 5D). Tumors from Patient_hi showed more immunosuppressive features in myeloid cells and reduced cytotoxicity in T cells in scRNA-seq data (Figure 5E). These findings were corroborated by immunofluorescence staining of patient tissues, which revealed an increased abundance of Ki-67 positive percentage, IBA1⁺CD206⁺ suppressive myeloid cells and a decreased presence of CD8⁺GZMB⁺ effector T cells in Patient_hi compared to Patient_lo (Figures 5F and S7F-S7H). In The Cancer Genome Atlas (TCGA) and Chinese Glioma Genome Atlas (CGGA) datasets, the Ribo-Transferʰⁱ gene signature derived from scRNA-seq differential expression analysis correlated with higher tumor grade and IDH-wildtype status in glioma (Figure S7I). Patients were further stratified into high- and low-scoring groups based on this Ribo-Transferʰⁱ signature. The high-scoring group exhibited elevated translational activity, enhanced proliferation, myeloid-derived immunosuppression and CD8⁺ T cell dysfunction features and was associated with a poorer prognosis (Figures 5G, S7J and S7K). Collectively, these results underscore the importance of ribosome transfer in driving GBM malignancy, indicating that interventions directed against this feature may provide clinical benefit.

**Figure 5.**
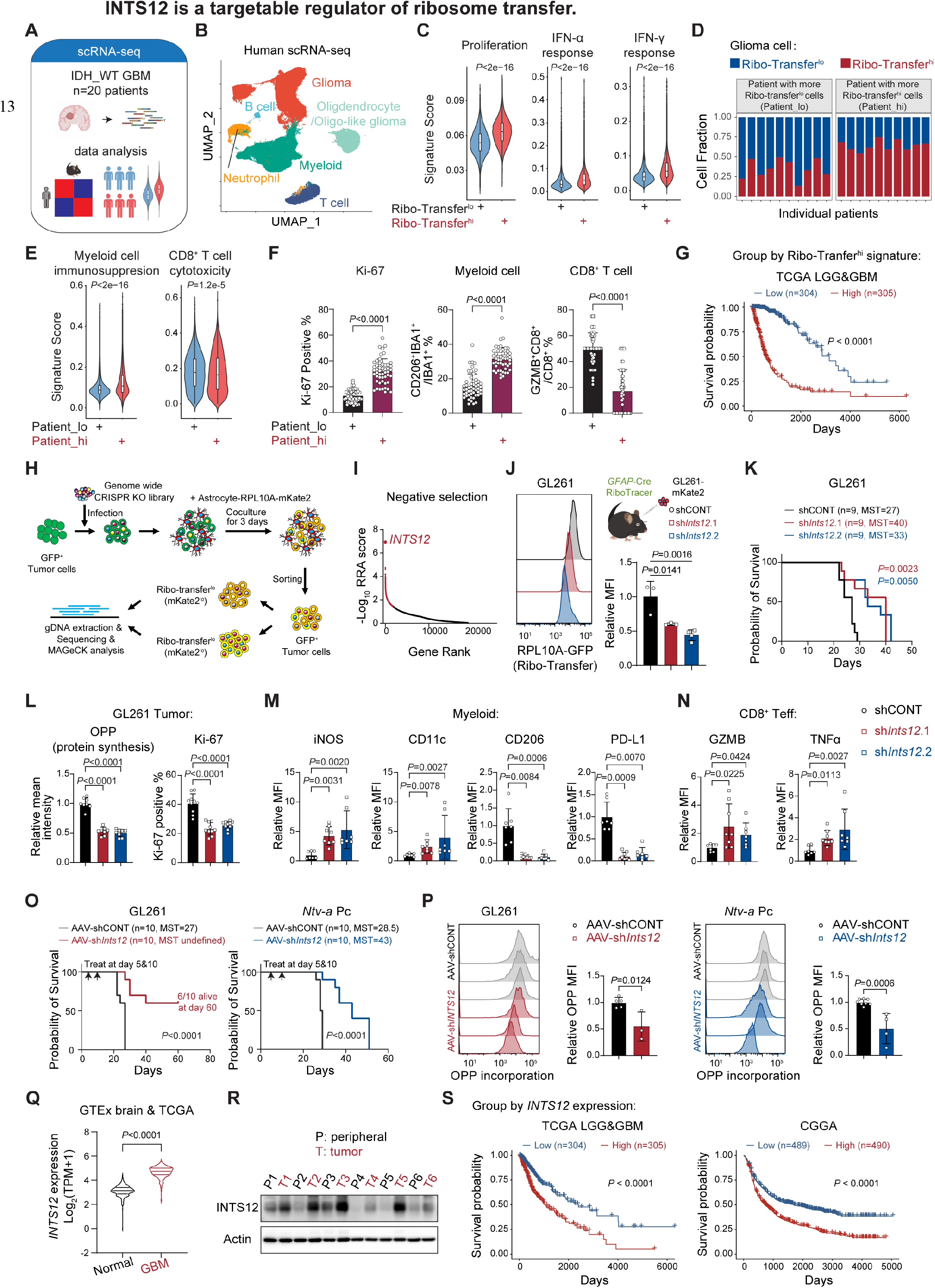
INTS12 is a targetable regulator of ribosome transfer. (A) Schematics of scRNA-seq of 20 IDH_WT GBM. (B) UMAP plot showing the identified cell clusters from (A). (C) Violin plots showing indicated signature scores in Ribo-Transfer^lo^ (n=37,779 cells) and Ribo- Transfer^hi^ (n=37,780 cells) GBM cells from scRNA-seq. (D) Fractions of Ribo-Transfer^lo^ and Ribo-Transfer^hi^ GBM cells in individual patients. (E) Violin plots showing immunosuppression score in myeloid cells (Patient_lo: n= 45,703 cells; Patient_hi: n= 27,713 cells) and cytotoxicity score in T cells (Patient_lo: n= 8,233 cells; Patient_hi: n= 9,471 cells) from scRNA-seq data. (F) Quantification of Ki-67^+^ percentage, suppressive myeloid cells (IBA1^+^CD206^+^), and effector T cells (CD8^+^GZMB^+^) in immunofluorescence-stained brain sections (n=10 patients per group, 5 fields per patient, 50 fields total per group). (G) Kaplan–Meier survival curves of patients stratified by low (n=304) vs. high (n=305) Ribo- Transfer^hi^ signature in the TCGA_LGG&GBM dataset. Cutoff, median ssGSEA signature score. (H) Schematics of a genome-wide CRISPR screen to identify drivers of astrocytic ribosome transfer in GBM cells. (I) Gene rank of negative selection (Ribo-Transfer^hi^ vs. Ribo-Transfer^lo^) in the CRISPR screen. Higher y-axis values indicate greater loss of ribosome transfer upon gene knockout. Red denotes significant hits. (J) Representative flow plots and quantification of RPL10A-GFP in *GFAP*-Cre; RiboTrancer mice implanted with GL261 with or without *Ints12* knockdown (shCONT: n=3; sh1: n=3; sh2: n=4). (K) Kaplan-Meier survival curves of immunocompetent mice implanted with GL261 with or without *Ints12* knockdown (n=9 per group). MST, median survival time. (L) Quantification of mean OPP fluorescence intensity (n=8 randomly selected fields from at least three mice) and Ki-67^+^ percentage (n=10 randomly selected fields from at least three mice) in brain sections from GL261-bearing mice with or without *Ints12* knockdown. (**M** and **N**) Quantification of indicated proteins in myeloid cells or CD8^+^ effectors T cells by flow cytometry in brain tumors initiated by GL261 with or without *Ints12* knockdown. (shCONT: n=7; sh1: n=8; sh2: n=7). (O) Kaplan–Meier survival curves of GL261- or *Ntv-a* Pc-bearing immunocompetent mice treated with vector or AAV-sh*Ints12* (n=10 per group). Arrows indicate the treatment timepoint (days 5 and 10). MST, median survival time. (P) Representative flow plots and quantification of OPP incorporation in brain tumors from GL261- or *Ntv-a* Pc-bearing immunocompetent mice treated with vector or AAV-sh*Ints12* (GL261: n=5 for vector, n=3 for AAV; *Ntv-a* Pc: n=8 for vector, n=4 for AAV). (Q) Violin plots showing *INTS12* expression in normal (n=295) and GBM (n=232) patients in GTEx brain and TCGA datasets. (R) Immunoblots showing INTS12 protein levels in paired GBM and peripheral tissues. (S) Kaplan-Meier survival curves of patients stratified by low vs. high *INTS12* expression in the TCGA_LGG&GBM (low: n=304; high: n=305) and CGGA (low: n=489; high: n=490) datasets. Cutoff, median. Data are represented as the mean ± SD in (F), (J), (L-N) and (P). Violin plots show the overall distribution of data points; boxes or horizontal lines indicate median, upper, and lower quartiles. Individual data points represent one mouse unless otherwise specified. Two-tailed Wilcoxon rank- sum test for (C) and (E). Two-tailed unpaired t test for (F), (P) and (Q). One-way ANOVA followed by multiple comparisons for (J) and (L). Kruskal-Wallis test followed by multiple comparisons for (M) and (N). Log-rank test for (G), (K) and (S). See also Figures S7-S9.

TNT-like structures serve as conduits for ribosome transfer as well as for the intercellular exchange of mitochondria and other cargoes^7^, rendering them a non-specific target and raising concerns about potential toxicity upon therapeutic intervention. To more precisely identify mediators of ribosome transfer and uncover targetable factors, we performed a genome-wide CRISPR knockout screen in patient-derived GBM cells (Figure 5H). After direct co-culture with astrocytes, we sorted Ribo-Transferʰⁱ and Ribo-Transferˡᵒ tumor cells and sequenced the integrated single guide RNAs. Genes whose knockout suppressed ribosome transfer (negative selection) were enriched in pathways related to cell communication and cell surface receptors (Figure S8A), with *INTS12* emerging as the top candidate (Figure 5I). INTS12 is a non-catalytic subunit of the integrator complex, an assembly of at least 19 subunits that binds RNA polymerase II and regulates gene transcription^50^. Previous studies have suggested context-dependent roles for INTS12 in cellular biology^51–53^. Knockdown of INTS12 in tumor cells significantly reduced ribosome uptake in the patient-derived coculture system and the murine reporter model (*GFAP*-Cre; RiboTracer) (Figures 5J and S8B-S8D). Loss of INTS12 in both murine and patient-derived orthotopic GBM models prolonged survival, reduced global protein synthesis and suppressed tumor cell proliferation (Figures 5K, 5L and S8E-S8H). Cell-autonomous INTS12 loss did not directly suppress tumor cell proliferation (Figures S8I-S8K). However, when these INTS12 knockdown tumor cells were co-cultured with astrocytes and then re-isolated, their proliferation was significantly inhibited (Figures S8L-S8N). These results indicate that the effect of INTS12 on GBM cell proliferation depends largely on their interactions with astrocytes and on the ribosome transfer it mediates. In immunocompetent mouse models, INTS12 knockdown led to an improved immune landscape within tumors, characterized by upregulation of activation markers (iNOS, CD11c) and downregulation of suppressive markers (CD206, PD-L1) on myeloid cells, and increased expression of GZMB and TNFα in CD8⁺ effector T cells (Figures 5M and 5N, gating strategy in S8O). Together, these findings suggest that INTS12 is a critical driver of ribosome acquisition in GBM cells, and that targeting INTS12 disrupts this process, alleviating the pro-proliferative and immunosuppressive tumor microenvironment.

To further evaluate the therapeutic potential of targeting INTS12, we performed intracranial implantation of GL261 cells and then intratumorally injected adeno-associated virus serotype DJ (AAV-DJ) expressing shRNAs. This treatment significantly prolonged mouse survival (Figure 5O). We also observed similar anti-tumor effects of INTS12 targeting in a genetically defined GBM model generated by primary tumor cells from *Ntv-a*; *Ink4a*^⁻/⁻^; *Pten*^f/f^ mice using the RCAS-PDGFB and *Nes*-Cre tv-a system (*Ntv-a* Pc) (Figure 5O), which has been reported to better recapitulate human GBM properties^54–56^. Analysis of brain tumors confirmed successful INTS12 knockdown within the malignant lesion, whereas INTS12 levels in the surrounding normal brain regions remained unchanged (Figures S8P and S8Q). OPP incorporation was markedly reduced (Figures 5P and S8P), indicating impaired global protein synthesis in tumor cells. To strengthen clinical relevance, we interrogated *INTS12* expression in clinical datasets. *INTS12* was significantly upregulated in GBM compared with normal brain tissue (Figures 5Q and 5R), and pan-cancer analysis revealed that GBM exhibited the highest *INTS12* expression among all tumor types (Figure S9A). *INTS12* expression positively correlated with both the Ribo-Transferʰⁱ signature and translational activity (Figure S9B). Patients with higher *INTS12* expressions showed elevated scores for proliferation, myeloid cell immunosuppression and CD8⁺ T cell dysfunction (Figure S9C). Higher *INTS12* expression was associated with poor prognosis in glioma (Figure 5S). Altogether, these findings support the rationale for targeting INTS12 as an emerging therapeutic strategy in GBM.

## DISCUSSION

Malignant progression is driven not only by cell-intrinsic genetic alterations but also by the exploitation of surrounding non-malignant cells^1^. In this study, we uncover an additional mode of intercellular communication: the direct transfer of functional ribosomes from astrocytes to GBM cells. This process supports protein synthesis in tumor cells, promotes their proliferation, and shapes an immunosuppressive microenvironment. The earliest clue for ribosome transfer came from the observation that Schwann cells, which tightly ensheath peripheral axons and transfer ribosomes to support local axonal translation upon nerve injury^57–59^. Recently, cell-to-cell ribosome transfer was observed to rescue protein synthesis in pancreatic cancer cells under stress *in vitro*^60^. However, whether ribosome transfer occurs *in vivo* within a complex heterotypic microenvironment, and how it contributes to tumor malignancy and orchestrates tumor immunity remains completely unknown. Using an *in vivo* ribosome-tracing model (RiboTracer) and *in vitro* co-culture systems, we provide high-level evidence that ribosome transfer takes place from astrocytes to GBM cells in both murine and patient-derived models. Moreover, transferred ribosomes are functional, accelerating tumor cell proliferation and reshaping tumor immunity to exacerbate GBM progression (Figure S9D, graphic abstract). Our findings deepen the understanding of ribosome transfer and add an additional layer of translational control and intercellular communication in cancer.

Beyond simply providing a quantitative boost to protein synthesis, astrocyte-derived ribosomes appear to carry an intrinsic translational preference. Using a cell-source-traceable ribosome profiling approach, we found that astrocyte-derived ribosomes preferentially translate transcripts involved in proliferation and IFN response. Under pathological conditions, astrocytes undergo reprogramming toward a reactive state, exhibiting context-dependent features that manifest as either protective or detrimental depending on the disease type and stage^17,18^. Proliferation is a commonly observed feature. Homeostatic astrocytes rarely divide, whereas reactive astrocytes can re-enter the cell cycle in response to brain injury, GBM, or brain metastasis^46,61–63^. IFN-responsive astrocytes have also been widely reported in neurological diseases, where they limit central nervous system inflammation in experimental autoimmune encephalomyelitis (EAE) models^19,64–66^. In the setting of GBM or brain metastasis, IFN-responsive astrocytes promote immunosuppression through upregulation of PD-L1 and secretion of pro-tumor cytokines^46–48^. Our observation that astrocyte-donated ribosomes preferentially translate proliferation- and IFN response-related transcripts mirrors the activated phenotype of tumor-associated astrocytes. These findings suggest that astrocytes within the GBM microenvironment not only serve as passive donors but also actively instruct the tumor’s translational program to fuel malignant progression.

The mechanistic basis for this translational selectivity remains an intriguing question.One possibility is that ribosomes are co-transferred with astrocyte-derived transcripts. However, given that the median half-life of mRNAs is approximately 7 hours^67^, it remains uncertain whether this mechanism alone can account for the sustained phenotypes in recipient tumor cells. Alternatively, astrocyte-derived ribosomes may carry distinct ribosomal protein compositions, co-factors, or post-translational modifications that confer preferential binding to specific mRNA subsets, as ribosomal components and modifications are commonly dynamic^68,69^. Further studies on ribosomal properties will elucidate how transferred ribosomes shape translation and whether this pathway can be therapeutically targeted.

We observed that astrocyte-to-tumor ribosome transfer depends on direct contact via TNT-like structures. Although ribosomes have been reported to exist extracellularly^70^, we did not detect significant transfer under indirect co-culture conditions, supporting a requirement for direct cell-to-cell contact. TNT are intercellular conduits that transfer mitochondria and various other cargoes^7^. In glioma, TNT-like structures *in vivo* appear thicker and longer and are also referred to as TMs^11^. Previous studies have shown that these structures enable functional intercellular networks among tumor cells and astrocytes, contributing to tumor invasion, recurrence, and cancer-neuron interactions^71–73^. Whether ribosome transfer through TNT-like structures similarly establishes a translational network and participates in these malignant behaviors remains to be investigated. Although TNT-like structures are open-ended, the transfer of certain cargoes can exhibit clear directionality. Using reciprocal labeling, we observed that ribosomes are predominantly acquired by tumor cells from astrocytes, a pattern consistent with the previously reported directionality of mitochondrial hijacking. The underlying mechanisms and the drivers governing this directional transfer warrant further study.

Through a genome-wide CRISPR knockout screen, we identified INTS12 as a critical and targetable driver of ribosome uptake in GBM. Genetic ablation of INTS12 in GBM cells reduced ribosome transfer, hindered tumor proliferation and remodeled the immune microenvironment. INTS12 is a component of the integrator complex, an assembly of at least 19 subunits that binds RNA polymerase II and cleaves nascent RNA transcripts, thus terminating non-productive transcription at protein-coding genes and processing diverse noncoding RNAs^50^. INTS12 is not a core catalytic subunit, and its structure has remained poorly resolved due to intrinsic flexibility^50,74^. Although the core subunits of the integrator complex are essential, loss of INTS12 appears to exert context-dependent functions. For instance, INTS12 knockdown suppresses proliferation in human bronchial epithelial cells^52^ but does not affect viability of CD4⁺ T cells^51^. A recent study revealed that INTS12 phosphorylation serves as a bridge linking UV-induced RNA damage signaling to DNA repair, and that INTS12 knockout exacerbates UV damage without affecting cell survival following ionizing radiation or calicheamicin treatment^53^. These observations underscore the context-specific nature of INTS12 functions. In this study, we found that INTS12 knockdown in GBM cells does not affect cell-autonomous proliferation yet attenuates tumor cell growth by orchestrating astrocytic ribosome transfer. Critically, pan-cancer analysis shows that INTS12 expression is highest in GBM among all cancer types and normal tissues, and correlates with poor survival in clinical datasets. This supports its potential as a compelling and specific therapeutic target that disrupts astrocyte-to-GBM ribosome transfer with minimal effects on normal tissues. Future studies are warranted to elucidate the mechanism by which INTS12 regulates ribosome transfer and to develop therapeutic strategies targeting this process, including T cell receptor-engineered T cell (TCR-T) therapy. We are actively pursuing this avenue and aim to advance toward clinical evaluation in the coming years.

In summary, we uncover astrocyte-to-GBM ribosome transfer as a driver of malignancy. Transferred ribosomes are functional, reconstructing the tumor translatome toward a proliferative and immunosuppressive state. We identify INTS12 as a key and targetable mediator of this process, whose loss impairs ribosome uptake and tumor progression, representing a promising therapeutic node that disrupts a non-cell-autonomous translational addiction in brain cancer.

## RESOURCE AVAILABILITY

### Lead contact

Requests for further information and resources should be directed to and will be fulfilled by the lead contact, Nu Zhang.

### Materials availability

This study did not generate new, unique reagents.

### Data and code availability

Raw sequencing data for murine scRNA-seq, ribosome-seq and matched bulk RNA-seq of Ribo-Transferˡᵒ and Ribo-Transferʰⁱ GL261 cells, as well as human scRNA-seq (20 GBM samples), ribosome-seq and matched bulk RNA-seq of IP-mKate2 and IP-Flag, and genome-wide CRISPR screen sequencing, have been deposited in the Genome Sequence Archive (GSA) and GSA-Human database at the National Genomics Data Center (NGDC) under accession IDs CRA045162, CRA045163 (mouse) and HRA019347, HRA019164 (human), respectively. All data in the main text or the supplementary materials are publicly available as of the date of publication. This paper does not report original code.

## ACKNOWLEDGMENTS

We sincerely appreciate all the patients who participated in the study. This work was funded by the National Natural Science Foundation of China (82125024; 82541020; 82341007; 82192894; 82573307; 82403655), Guangdong Major Project of Basic Research (2025B0303000013), Guangdong S&T Program (2025B0101130004), Guangdong Basic and Applied Basic Research Foundation (2025B1515020015), the Science and Technology Planning Project of Guangdong Province (2023B1212060018).

## AUTHOR CONTRIBUTIONS

Conceptualization: X.W., N.Z.; methodology: X.W., C.Z., L.Chen, J.Chen; investigation: X.W., C.Z., L.Chen, J.Chen, Y.Z., L.Cheng, Y.P., X.Z., M.G., F.L., Y.L., J.Cheng, Y.G., J.Z., X.Y., N.H.; visualization: X.W., C.Z., L.Chen; funding acquisition: N.Z., X.W.; project administration: N.Z., F.X.; supervision: N.Z., F.B., X.L.; writing-original draft: X.W., C.Z., L.Chen; writing-review & editing: N.Z..

## METHODS

### EXPERIMENTAL MODEL AND STUDY PARTICIPANT DETAILS

#### Human specimens

This study utilized pathologically verified GBM and peritumoral tissues obtained from surgical resections at the Department of Neurosurgery, First Affiliated Hospital of Sun Yat-sen University. Written informed consent was obtained from all patients prior to sample collection. Patient information for the samples subjected to scRNA-seq is provided in Table S1. The research was approved by the Clinical Research Ethics Committee of the First Affiliated Hospital of Sun Yat-sen University ([2020]322) and adhered to the guidelines of the Declaration of Helsinki.

#### Mouse strains and housing

C57BL/6J mice and BALB/c-nu mice were purchased from the Laboratory Animal Centre of the First Affiliated Hospital of Sun Yat-sen University. RiboTracer (R26-CAG-LSL-Avi-GFP-*Rpl10a*) mice (#NM-KI-210142) and R26-CAG-tdTomato mice (#NMX-KI-232404) were purchased from Shanghai Model Organisms Center, Inc. For pervasive cell labeling, *UBC*-Cre-ERT2 mice (#007179, The Jackson Laboratory) were crossed with RiboTracer mice, and the resulting offspring received intraperitoneal injections of tamoxifen (#T5648, Sigma-Aldrich) at 75 mg/kg once every 24 hours for three consecutive days, followed by a three-day waiting period, to generate *UBC*-Cre; RiboTracer mice. To label astrocytes, *GFAP*-Cre mice (#004600, The Jackson Laboratory) on a C57BL/6 background were purchased from Shanghai Model Organisms Center, Inc. and crossed with RiboTracer mice to generate *GFAP*-Cre; RiboTracer. Mice were housed under specific-pathogen-free conditions at an ambient temperature of 20-26 °C, relative humidity of 30-70%, and a 12-hour light/dark cycle prior to experimental use. Both male and female animals were included in the study. All mouse experiments were conducted in accordance with the Clinical Research and Laboratory Animal Ethics Committee of the First Affiliated Hospital of Sun Yat-sen University ([2023]132).

#### Cell derivation and culture

GL261 cells were purchased from EK Biosciences (#CC-Y2133). CT2A cells were purchased from Millipore (#SCC194). The 293T line was purchased from ATCC (#CRL-3216). Primary human normal astrocytes were purchased from Lonza (#CC-2565). Above cells were cultured in DMEM medium (#11995065, Gibco) supplemented with 10% fetal bovine serum (FBS; #MK1123-500C, MIKX). *Ntv-a* Pc primary GBM cells were derived from brain tumors induced by intracranial injection of RCAS-PDGFB and RCAS-Cre retrovirus into *Ntv-a*; *Ink4a*^⁻/⁻^; *Pten*^f/f^ mice, as previously reported by our group^75^. Patient-derived #147 GBM cells were established in our laboratory from a GBM specimen of a 68-year-old female patient. Patient-derived #23 GBM cells were established from a recurrent GBM biopsy obtained from a 63-year-old male patient and were generously provided by Prof. Xiuxing Wang (Nanjing Medical University). *Ntv-a* Pc and patient-derived cells were cultured in Neurobasal media (#21103049; Gibco) supplemented with B27 without vitamin A (#12587010; Gibco), 20 ng/ml recombinant human EGF (#236-EG-01M; R&D Systems), and 20 ng/ml recombinant human bFGF (#4114-TC-01M; R&D Systems), sodium pyruvate (#11360070; Gibco), GlutaMAX (#35050061; Gibco) and streptomycin/penicillin (#15140122; Gibco). For direct coculture experiments, donor and recipient cells were seeded at a 1:1 ratio onto Matrigel-coated plates and cocultured in serum-free complete Neurobasal media for 3 days. For inhibitor treatment, ML-141 (10 μM; #HY-12755, MedChemExpress) or L-778123 (10 μM; #HY-16273A, MedChemExpress) was directly added into the coculture system for 3 days, with an equal volume of DMSO as vehicle control. For indirect coculture, donor cells were seeded on transwell inserts (0.4-μm pore size) and recipient cells were seeded on the lower chamber. All cultures were maintained at 37 °C in a humidified atmosphere containing 20% O₂ and 5% CO₂. Short tandem repeat profiling was conducted annually to verify the identity of each tumor model used in this study. Mycoplasma contamination was routinely tested via PCR analysis of culture supernatants.

#### Orthotopic GBM models and *in vivo* treatment

Syngeneic orthotopic GBM models were established by intracranial injection of tumor cells into the right cerebral cortex of C57BL/6J mice at a depth of 3.5 mm following standard procedures. The injection volumes contained 200,000 GL261 cells, 300,000 CT2A cells, or 300,000 *Ntv-a* Pc primary GBM cells. Patient-derived xenograft models were established by intracranial injection of 100,000 #147 or #23 GBM cells into the same brain region of BALB/c-nu mice. For genetically ablation of INTS12 *in vivo* via AAV treatment, recombinant AAV-DJ carrying a plasmid expressing four different shRNAs targeting *Ints12* (sequences provided in Table S2) or non-targeting shRNAs (generated by WZ Biosciences) was intratumorally injected at a dose of approximately 2.5 × 10^11^ viral particles per injection on days 5 and 10 after tumor cell implantation. All housing conditions and animal health were monitored by a licensed veterinarian. Mice were observed daily and euthanized upon the onset of neurological symptoms, including hunched posture, altered gait, lethargy, and weight loss. Brains were collected postmortem, fixed in 4% paraformaldehyde, and processed for paraffin embedding. Histological analysis was performed on tissue sections using hematoxylin and eosin (HE) staining. Survival data were analyzed using GraphPad Prism software (v9.5.0), and statistical significance was determined by log-rank test.

## METHOD DETAILS

### Isolation of primary cells from *UBC*-Cre-ERT2 RiboTracer mice and coculture

Brains from neonatal *UBC*-Cre-ERT2; RiboTracer mice (<24 h old) were rapidly collected for the isolation of primary astrocytes, microglia, and neurons, following previously described procedures^76,77^ with modifications. For astrocyte and microglia isolation, cerebral cortices were dissected using ophthalmic forceps, and the meninges were carefully removed under a stereomicroscope. The tissue was minced in prechilled serum-free DMEM/F12 medium (#C11330500BT, Gibco) and then digested with 0.125% trypsin (#25200072, Gibco) at 37 °C with shaking at 150 rpm for 15 min. Digestion was stopped by adding an equal volume of DMEM containing 10% fetal bovine serum (FBS) and 1% penicillin/streptomycin. The cell suspension was centrifuged at 500 × g for 5 min, the supernatant was discarded, and the pellet was resuspended in DMEM with 10% FBS and 1% penicillin/streptomycin. The suspension was passed through a 70-μm cell strainer, and the cell concentration was adjusted before seeding onto a T25 culture flask pre-coated with 0.1 mg/mL poly-L-lysine (#P6407, Sigma-Aldrich). After overnight culture, non-adherent cells were removed by medium aspiration and PBS washing, followed by replacement with fresh complete DMEM. Mix cultures were maintained for 7 days with medium replacement every 2-3 days to reach confluency. Oligodendrocyte progenitor cells (OPCs) were removed by gentle tapping, repeated several times. The flasks were then shaken at 37 °C and 200 rpm. After shaking for 3 h, the supernatant containing primary microglia was collected and centrifuged at 1200 rpm for 5 min. The microglia were resuspended in complete DMEM medium for subsequent coculture. The remaining adherent cells in the T25 flasks were enriched in primary astrocytes. Both primary microglia and astrocytes were cultured for an additional 48 h for stabilization before subsequent experiments. To obtain primary neurons, hippocampi were dissected under a stereomicroscope and transferred into dissection buffer (phenol red-free high-glucose DMEM (#21063029, Gibco) supplemented with 1% horse serum (#16050130, Gibco) and 1% penicillin/streptomycin). Residual meninges and blood vessels were removed as completely as possible. The tissue was gently triturated in fresh dissection buffer, minced, and resuspended in digestion buffer containing 1 mg/mL papain (#P4762-100MG, Sigma-Aldrich) and 1 mg/mL DNase I (#4536282001, Sigma-Aldrich) in phenol red-free DMEM. The suspension was transferred to a 15-mL centrifuge tube and incubated at 37 °C with gentle trituration every 10 min. After 30 min, digestion was terminated by adding an equal volume of DMEM containing 10% fetal bovine serum (FBS) and 1% penicillin/streptomycin. The cell suspension was centrifuged at 700 rpm for 2 min, and the supernatant was discarded. The pellet was resuspended in 2 mL DMEM containing 1 mg/mL DNase I, gently triturated, and allowed to settle on ice for 3 min. The uppermost 1 mL of the suspension was transferred to a new tube. An additional 1 mL of DMEM with 1 mg/mL DNase I was added to the remaining suspension, which was again allowed to settle on ice, and the uppermost 1 mL was collected. This process was repeated three times. The collected cell suspension was centrifuged at 4 °C and 1000 rpm for 5 min. After removal of the supernatant, cells were resuspended in an appropriate volume of DMEM with 10% FBS and 1% penicillin/streptomycin and filtered through a 70-μm cell strainer. Cell density was adjusted with the same medium before seeding onto T25 culture flasks pre-coated with 0.1 mg/mL poly-L-lysine. After incubation at 37 °C for 2 h, the medium was removed and replaced with neuronal culture medium consisting of Neurobasal-A (#10888022, Gibco) supplemented with 2% B27, 5% horse serum, 1% GlutaMAX, and 0.5% penicillin/streptomycin. Half of the medium was replaced every 3-4 days. Neurons were used for subsequent experiments after developing typical neuronal morphology, usually after about 7 days in culture.

Bone marrow derived macrophages (BMDMs) were isolated from femurs and tibiae of adult *UBC*- Cre-ERT2; RiboTracer mice following a previously reported protocol^78^. After euthanasia, the femurs and tibiae were carefully excised, and the surrounding skin and muscle tissue were removed without damaging the bones. The cleaned bones were immersed in 75% ethanol for 5 min for sterilization, then transferred to PBS to remove residual ethanol. Both ends of each bone were cut with ophthalmic scissors, and the bone marrow cavity was flushed with complete RPMI 1640 medium (#C11875500BT, Gibco) using a 1-mL syringe. The flushing procedure was repeated three times until the bones became visibly pale. The collected marrow suspension was gently pipetted to disperse clumps and filtered through a 70-μm cell strainer. The filtrate was centrifuged at 1500 rpm for 5 min. The supernatant was discarded, and the cell pellet was resuspended in red blood cell lysis buffer (#00-4333-57, eBioscience) for 3 min at room temperature. Lysis was stopped by adding 10 mL PBS, followed by centrifugation at 1500 rpm for 5 min. The resulting cell pellet was resuspended in complete RPMI 1640 medium supplemented with 10 ng/mL M-CSF (#Z03275-50, GenScript) and seeded into T25 culture flasks. Cells were maintained at 37 °C in a humidified incubator with 5% CO_2_. On day 3, half of the culture medium was replaced. On day 5, the medium was completely replaced. On day 7, BMDMs were fully differentiated and ready for downstream assays.

Upon reaching maturity and being ready for experiments, primary cells were treated with 1 μM 4-hydroxytamoxifen (4-OHT, #S8956, Selleck) for 3 days. The medium was then replaced with fresh culture medium, and cells were cultured for an additional 3 days to allow Cre-mediated recombination, resulting in GFP labeling of ribosomes in primary cells for subsequent coculture experiments. To avoid the loss of cell viability caused by enzymatic dissociation, tumor cells were directly added to primary cell cultures at a 1:1 ratio (based on the initial seeding density) using a 1:1 mixture of their respective culture media. After 3 days of coculture, cells were collected for flow cytometric analysis to assess ribosome transfer.

### Immunofluorescence staining

Cells were plated on Matrigel-coated coverslips and fixed with 4% paraformaldehyde. Paraffin-embedded sections were deparaffinized and underwent antigen retrieval in sodium citrate buffer in a microwave oven for 15 min. Coverslips and sections were permeabilized with 0.3% Triton X- 100, blocked with 5% bovine serum albumin (BSA), and incubated with primary antibody at 4 °C overnight and appropriate secondary antibody for 2 h at RT. Nuclei were counterstained with 4′,6-diamidino-2-phenylindole (DAPI). Antibodies used in this study are provided in Table S3. F- actin was labeled by fluorescence-conjugated phalloidin (Beyotime). Images were acquired using an Olympus BX63 fluorescence microscope, a Zeiss LSM 880 confocal microscope with Airyscan detector, or a Nikon N-SIM S S Super Resolution Microscope. Z-stack images were reconstructed into 3D renderings using Imaris software (version 9.01).

### Isolation of single cells

Mouse brain tumors were obtained after administration of a lethal dose of anesthetics followed by transcardial perfusion with PBS. The tumors were dissected and rinsed thoroughly in PBS to remove surface blood contamination. The tissue was then minced into small pieces and digested in DMEM containing 10% FBS, 1 mg/mL collagenase type IV (#17104015, Gibco), 0.1 mg/mL hyaluronidase (#H8030, Solarbio), and 0.1 mg/mL DNase I (#HY-108882, MedChemExpress) at 37 °C for 30 min, with gentle trituration every 10 min. Following digestion, the tissue suspension was passed through a 70 µm strainer to remove clumps and obtain single cell suspensions. After centrifugation at 500 × g for 5 min and removal of the supernatant, the cell suspension was subjected to density gradient centrifugation using a 30% Percoll (#17089102, Cytiva) solution at 500 × g with the lowest brake for 20 min at 4 °C to remove the myelin-rich upper layer. The remaining cell pellet was resuspended in RBC lysis buffer (#AR1118, Boster) for 3 min at room temperature. RBC lysis was terminated by adding 10 mL PBS, and cells were collected by centrifugation at 200 × g for 3 min. The supernatant was discarded, and the cells were resuspended in staining buffer (PBS with 2% FBS) for downstream applications, such as flow cytometry, fluorescence-activated cell sorting (FACS), or scRNA-seq.

For human brain tumors, tissues were transported on ice in a sterile culture dish containing 10 mL of 1x Dulbecco’s Phosphate-Buffered Saline (DPBS; #14190144, Thermo Fisher) to remove residual storage solution and then minced on ice. Tissue fragments were digested in 0.25% trypsin and 10 μg/mL DNase I dissolved in PBS containing 5% FBS. Digestion was carried out at 37 °C with shaking at 50 rpm for approximately 40 min. To increase cell yield and viability, dissociated cells were repeatedly collected at 20-min intervals during digestion. The resulting cell suspension was filtered through a 40-μm cell strainer, and red blood cells were removed using 1x Red Blood Cell Lysis Solution (#00-4333-57, Thermo Fisher). Dissociated cells were then washed with 1× DPBS containing 2% FBS. Cells were assessed for viability and then processed for scRNA-seq.

### Flow cytometry analysis

Cell suspensions were first labeled with surface fluorescent antibodies on ice for 30 min, then fixed and permeabilized using the True-Nuclear™ Transcription Factor Buffer Set (#424401, BioLegend), followed by staining for intracellular markers. Antibodies used in this study are listed in Table S3. Dead cells were labeled with Fixable Viability Dye eFluor 455UV (#65-0868-14, eBioscience) and excluded. Gating strategies are provided in the supplemental figures. Flow cytometry was performed on a Cytek Aurora flow cytometer or a BD FACSymphony A3 flow cytometer, and the data were analyzed using FlowJo software (version 10.6.2).

### Fluorescence-activated cell sorting (FACS)

FACS was carried out on a BD FACSAria Fusion flow cytometer. GFP fluorescence was excited with a 488 nm laser and detected through a 530/30 nm filter. mKate2 fluorescence was excited with a 561 nm laser and detected through a 610/20 nm filter. Sorting strategies are provided in the corresponding figures.

### Immunoblotting

Cells were lysed in RIPA buffer (50 mM Tris pH 7.4, 150 mM NaCl, 1% NP-40, 0.5% sodium deoxycholate, 0.1% SDS) containing protease inhibitor cocktail. For immunoblotting, 30-50 μg proteins were subjected to SDS-PAGE and then transferred to polyvinylidene difluoride (PVDF) membrane (#1620177, Bio-Rad). Membranes were blocked with 5% skim milk, incubated with primary antibodies, followed by appropriate horseradish peroxidase-conjugated secondary antibodies. Protein signals were visualized using ECL substrate. Antibodies used in this study are listed in Table S3.

### Reverse transcription and quantitative polymerase chain reaction (RT-qPCR)

Total RNA was isolated and complementary DNA (cDNA) was synthesized using Evo M-MLV RT Master Mix (#AG11706, Accurate Biology). qPCR was performed using SYBR Green Pro Taq HS Premix (#AG11718, Accurate Biology) in a QuantStudio 5 system (Applied Biosystems). Primer sequences are detailed in Table S2.

**Protein synthesis measurement *in vivo* and *in vitro***

To measure protein synthesis *in vivo*, GBM-bearing mice were intracranially injected with 20 μl of 1.5 mM *O*-Propargyl-Puromycin (OPP; HY-15680, MedChemExpress) into the contralateral hemisphere using the following coordinates: −0.5, 1.0, −2.5 mm. Mice were sacrificed 30 min later, and brains were harvested for for either cell isolation followed by flow cytometry or tissue sections for immunofluorescence staining. OPP incorporation signals were detected by click reaction using reagents from the BeyoClick™ EdU Cell Proliferation Kit with AF488 or AF647 (#C0071S/C0081S, Beyotime). For *in vitro* protein synthesis assessment, cells were treated with 1 μM puromycin for 30 min followed by immunoblotting with an anti-puromycin antibody, or with 15 μM OPP for 30 min following by click reaction.

### Plasmids and lentiviral infection

Lentiviral expression vectors encoding RPL10A-mKate2, RPL36-mKate2, RPL10A-Flag, RPL36-Flag, GFP, or mKate2, as well as lentiviral clones expressing shRNAs targeting human *TNFAIP2*, human *INTS12*, murine *Ints12*, or control sequences, were generated by Wuhan Miaoling Biotechnology. Lentiviral particles were produced by co-transfecting 293T cells with the respective constructs and the packaging plasmids pMD2.G and psPAX2 using PEI. Viral supernatants were collected 48 h post-transfection and concentrated by precipitation with PEG8000. Cells were infected with lentivirus for 24 h, followed by a 24-h recovery period before antibiotic selection. For cells intended for OPP or puromycin incorporation assays, selection markers were restricted to blasticidin or neomycin, as puromycin resistance was avoided to prevent potential interference with protein translation measurements.

### Cell proliferation and EdU incorporation assay

Cell viability was assessed by seeding 7,500 cells per well into Matrigel-coated 96-well plates. At the indicated time points, cell viability was measured using the Cell Counting Kit-8 (#C0038, Beyotime). Absorbance values at 450 nm were normalized to day 1 measurement. For direct cell counting, 1x10^5^ cells were seeded into six-well plates, and viable cells were counted using Trypan blue staining. Counts were normalized to the day 1 value. EdU incorporation assay was performed using the BeyoClick™ EdU Cell Proliferation Kit (Beyotime) according to the manufacturer’s instructions. Cells were incubated with 10 μM EdU for 2 hours, followed by click reaction and imaging. The proportion of EdU-positive cells was determined using ImageJ software by calculating the ratio of EdU-positive nuclei to total DAPI-stained nuclei.

### Single-cell RNA sequencing (scRNA-seq)

Single-cell suspensions of brain tumors were prepared as described in the "Isolation of single cells" section. For murine samples, brain tumors initiated by GL261-mKate2 cells in *GFAP*-Cre; RiboTracer mice were dissociated into single cells. Viable CD45^-^ mKate2^+^ cells were further sorted into Ribo-Transferˡᵒ (GFPˡᵒ) and Ribo-Transferʰⁱ (GFPʰⁱ) populations at a 40% cutoff for each (i.e., the lowest and highest 40% of GFP signal intensity), which were then processed for library construction. For human GBM samples, no cell sorting was performed. The 10x Genomics platform was used for cDNA generation, followed by single-cell library construction and sequencing on the Illumina NovaSeq 6000 platform.

### scRNA-seq data analysis

#### Preprocessing and quality control of the scRNA-seq data

Raw sequencing data generated from the 10x Genomics platform were demultiplexed into FASTQ files using CellRanger mkfastq (v5.0.0). Sequencing reads were then aligned against the reference genomes GRCh38-1.2.0 (human) and mm10-2020-A (mouse) via CellRanger count (v5.0.0). To integrate multi-run datasets, outputs were aggregated with CellRanger aggr (v5.0.0). For downstream analysis, scRNA-seq data processing and visualization were implemented in R (v4.5.3) using the Seurat package (v5.5.0)^79^. Quality control filtering was applied independently to each sample category with the following thresholds: (1) mouse GBM samples: maximum percentage of mitochondrial reads=10%, maximum number of UMIs=45,000, maximum number of genes=7,500, and minimum number of gene=500; (2) human GBM samples: maximum percentage of mitochondrial reads=10%, minimum number of gene=500. Doublet scoring was performed using the R package scDblFinder (v1.24.10)^80^ with default parameters to remove putative doublets from each sample. To further ensure the robustness and biological validity of downstream analyses, we manually curated the unsupervised clustering results for the human GBM data. Clusters that lacked biological reproducibility (i.e., being highly sample-specific or batch-driven) or exhibited mixed transcriptional profiles, such as co-expression of mutually exclusive lineage markers, were considered unresolved multiplets, low-quality cells, or droplets heavily contaminated by ambient RNA. These clusters represented approximately 0.56% of total cells and were considered technical artifacts, thus were excluded from further analyses. After their removal, the remaining high-quality cells were re-scaled and re-clustered. This process yielded 52,556 high-quality cells in the mouse GBM samples and 189,319 high-quality cells in the human GBM samples.

#### Data normalization, dimensionality reduction and clustering

For mouse GBM samples, normalization and variance stabilization were performed using the SCTransform function in Seurat (v5.5.0). For human GBM samples, due to the large dataset size that precluded the computational demands of SCTransform, data were first normalized using NormalizeData function, followed by feature selection via FindVariableFeatures function (vst method, top 2,000 highly variable genes) and scaling with ScaleData function. Principal component analysis (PCA) was then performed on the scaled variable genes, and the first 40 principal components (PCs) were retained. To correct for batch effects across samples within each dataset, the Seurat objects were integrated using Harmony (v2.0.3)^81^, and the corrected Harmony embeddings were used for downstream analysis. For each dataset, a shared nearest neighbor (SNN) graph was constructed using the Harmony dimensions (FindNeighbors, reduction = "harmony"), followed by graph-based clustering with the Louvain algorithm (FindClusters, resolution adjusted per dataset to yield stable cluster separation). Clusters were visualized using UMAP or *t*-SNE.

#### Cell type annotation

To distinguish individual cell types, we employed a three-step approach integrating unbiased clustering, copy number variation (CNV) inference, and expression of cell-type-specific marker genes. First, we used UMAP to visualize all cells in the same transcriptional space. Second, cells were classified as immune or non-immune based on *PTPRC* (encoding CD45) expression. Within the non-immune fraction, malignant tumor cells were identified by single-cell CNV inference using R package copyKAT (v1.1.0)^82^. Because oligodendrocytes and oligo-like glioma cells are difficult to distinguish based solely on marker expression in human GBM samples, we extracted CNV-positive cells as tumor cells when they resided within the same cluster. Finally, cell-type annotations were validated by examining the expression of canonical marker genes. For mouse GBM samples, the following markers were used: *Ptprc* for immune cells; *Aif1*, *Csf1r*, *Cd68* for myeloid cells; *Flt1*, *Esam*, *Pecam1* for endothelial cells; and *Fbln1*, *Col1a1*, *Col1a2* for fibroblasts. For human GBM samples, markers included: *PTPRC* for immune cells; *CX3CR1*, *P2RY12*, *CSF1R*, *CD163* for myeloid cells; *CSF3R*, *MPO* for neutrophils; *CD3E*, *CD3D* for T cells; *CD79A*, *MS4A1* for B cells; and *MOG*, *PLP1* for oligodendrocytes. Dot plots were generated using the top cluster-specific genes for each cell type.

#### Signature score analysis based on gene sets

To assess the status of cell clusters, we quantified transcriptional profiles using curated gene sets derived from prior studies^83,84^ and published database (e.g. Gene Ontology (GO) database). Signature scores were computed using the AddModuleScore_UCell function from the R package UCell (v2.12.0)^85^, which implements a rank-based statistic that is robust to dataset size and heterogeneity. The gene signatures used in this study are listed in Table S4.

#### Differential expression analysis and pathway enrichment

Differentially expressed genes (DEGs) between RiboTransfer^lo^ and RiboTransfer^hi^ cells were identified using the FindMarkers function in Seurat (v5.5.0) with the Wilcoxon rank-sum test. Genes were considered statistically significant if they met the following criteria: an absolute log₂ fold change > 0.25, an adjusted *P*-value < 0.05 (Bonferroni correction) and a minimum percentage of expression in at least 10% of cells (min.pct = 0.1). To investigate functional pathways associated with the identified DEGs, GO enrichment analysis was conducted using the R package clusterProfiler (v4.20.0)^86^. Pathways with a Benjamini-Hochberg corrected *P*-value < 0.05 were considered significantly enriched.

#### Mapping GBM cells onto cell states

To classify GBM cells into four molecular subtypes (MES-like, AC-like, OPC-like, NPC-like), we used AddModuleScore function in Seurat (v5.5.0) to calculate module scores based on previously published cell-state-specific gene sets^39^. The MES score was calculated as the average of MES1 and MES2, and the NPC score as the average of NPC1 and NPC2; these were then combined with the AC and OPC scores and standardized. Cells were mapped onto a four-quadrant system, where the Y-axis (*D* = max (OPC score, NPC score) - max (AC score, MES score)) distinguished OPC/NPC-positive from AC/MES-positive cells, and the X-axis represented relative differences between scores (OPC vs. NPC or AC vs. MES). Density plots were generated for Ribo-Transferʰⁱ and Ribo-Transferˡᵒ subpopulations using kernel density estimation, demarcating cell-state-specific distributions.

#### Classification of Ribo-Transfer^hi^ and Ribo-Transfer^lo^ human GBM cells

Human GBM cells were classified into Ribo-Transferʰⁱ and Ribo-Transferˡᵒ subpopulations using a predictive framework based on mouse GBM cells. The reference dataset consisted of scRNA-seq profiles from mouse GBM cells sorted into Ribo-Transferʰⁱ and Ribo-Transferˡᵒ groups via FACS. Using the SingleR algorithm (v2.12.0)^87^, transcriptional profiles of human GBM cells were compared against the annotated mouse reference dataset. A similarity score was calculated for each human cell, reflecting its transcriptional resemblance to mouse Ribo-Transferʰⁱ cells. Human GBM cells were then stratified into Ribo-Transferʰⁱ and Ribo-Transferˡᵒ groups based on whether their scores were above or below the median across all human GBM cells. This cross-species approach allowed for the functional stratification of human GBM cells based on ribosome transfer status.

### Ribosome profiling (Ribo-seq) and matched RNA-seq

For collection of Ribo-Transferˡᵒ and Ribo-Transferʰⁱ GL261 cells, these populations were isolated by FACS from tumor-bearing *GFAP*-Cre; RiboTracer mice as described above. The cell pellets were snap-frozen in liquid nitrogen and subjected to Ribo-seq and matched RNA-seq. For cell-source-traceable ribosome-seq, astrocytes expressed RPL10A-mKate2 to mark astrocytic ribosomes, and tumor cells expressed RPL10A-Flag to label their own ribosomes. After three days of direct coculture, GFP^+^ tumor cells were sorted by FACS. Cell pellets were snap-frozen in liquid nitrogen and then processed using a modified RiboTag protocol^88^. Briefly, cell pellets were resuspended in 950 μL ice-cold lysis buffer (50 mM Tris-HCl pH 7.4, 100 mM KCl, 12 mM MgCl_2,_ 1% NP-40, 1 mM DTT, 200 U/mL RNasin, 1 mg/mL heparin, 100 μg/mL cycloheximide (CHX), and 1x protease inhibitor). Lysates were incubated on ice for 30 min with gentle pipetting every 10 min, then centrifuged at 10,000 × g for 10 min at 4 °C. The supernatant was split into three aliquots corresponding to IgG control, Flag-IP, and mKate2-IP, and each was brought to 400 μL with lysis buffer. Antibodies (5 μg IgG, 5 μg anti-Flag, 5 μg anti-mKate2) and 2 μL RNasin (200 U/mL final) were added, and samples were rotated at 4 °C for 4 h. Meanwhile, 60 μL of Protein A/G magnetic beads per condition were washed with lysis buffer and rotated at 4 °C. After the 4-h incubation, beads were added to each lysate and rotated overnight at 4 °C. Beads were then collected on a magnetic rack, washed four times with high-salt buffer (50 mM Tris-HCl pH 7.4, 300 mM KCl, 12 mM MgCl_2_, 1% NP-40, 0.5 mM DTT, 100 μg/mL CHX), each wash rotating for 5 min at 4 °C. After the final wash, beads were snap-frozen in liquid nitrogen and stored at -80 °C until Ribo-seq and matched RNA-seq library preparation.

Ribo-seq and matched RNA-seq was conducted by Hangzhou NeoRibo Biotechnology Co., Ltd. The workflow is based on an advanced ribosome footprinting protocol that enables high-resolution mapping of ribosome-protected mRNA fragments, as previously described^89^. In brief, cells were lysed under conditions that preserve polysomes, and ribosome-mRNA complexes were digested with RNase I to generate ribosome-protected fragments (RPFs). The fragments were then purified, dephosphorylated, ligated to adaptors, and converted into cDNA libraries according to the QEZ- seq protocol. Matched RNA-seq libraries were prepared from the same cell lysates for transcriptome analysis. The constructed libraries were subjected to high-throughput sequencing on the Illumina NovaSeq X Plus platform using a paired-end 150 bp (PE150) strategy. This sequencing strategy, originally designed for small RNA libraries, ensures sufficient coverage and compatibility with ribosome-protected fragment profiling.

### Ribo-seq data analysis

For Ribo-seq, raw reads were first trimmed using cutadapt (v5.2)^90^ with the parameters -j 20 -- trim-n --match-read-wildcards -u 3 -n 4 -a AGATCGGAAGAGCACACGTCTG -a AAAAAAAA -a GAACTCCAGTCAC. The trimmed reads were sequentially mapped to mouse or human rRNA sequences (mm10/hg38) using Bowtie (v1.3.1)^91^ with parameters -seedlen=23, and rRNA-aligned reads were discarded. The remaining reads were aligned to the transcriptome (mm10/hg38) using STAR (v2.7.11b)^92^ with the parameters -outFilterMismatchNmax 2 -outFilterMultimapNmax 20 -outFilterMatchNmin 16 -alignEndsType EndToEnd. Downstream analyses were performed using the RiboWaltz package (v2.0)^93^ in R, which enabled quality assessment of ribosome footprints, including P-site offset calculation and evaluation of three-nucleotide periodicity. SAMtools (v1.21)^94^ was used for sorting, indexing and format conversion from SAM files. Raw ribosome footprints were quantified as the number of reads aligned to the CDS region for each gene using FeatureCounts (v2.0)^95^. For RNA-seq, raw reads were trimmed using cutadapt (v5.2). The cleaned reads were aligned to the mouse or human reference genome (mm10/hg38) using STAR (v2.7.11b) with default parameters. SAMtools (v1.21) was used for sorting, indexing and format conversion from SAM files. Raw counts were determined as the number of reads aligned to the exon of each gene using FeatureCounts (v2.0). Only uniquely mapped reads were considered for count assignment. Translational efficiencies were determined as the ratio of (normalized abundance determined by Ribo-seq)/(normalized abundance determined by RNA-seq). Different translational efficiencies were estimated using DESeq2 package (v1.49.4)^96^. Genes with |log_2_ fold changes|>1 and adjusted *P*-value<0.05 were regarded as significant. Downstream pathway enrichment was performed using gProfiler^97^ (https://biit.cs.ut.ee/gprofiler/). GSEA analysis was performed using ClusterProfiler (v4.20.0).

### Genome-wide CRISPR knockout screen and data analysis

A genome-wide CRISPR knockout library targeting 19,050 human coding genes and 1,864 miRNAs, comprising 65,383 single-guide RNA (sgRNA) constructs, was constructed using the lentiCRISPR v2 backbone (#1000000048, Addgene). The lentiviral pooled library was transduced into #147-GFP GBM cells at a multiplicity of infection (MOI) of 0.3, achieving a coverage of more than 500-fold with two replicates. Two days after transduction, puromycin was added at 2 μg/mL for selection. Five days post-transduction, the cells were directly co-cultured with human astrocytes expressing RPL10A-mKate2 at a 1:1 ratio for three days. RiboTransferlo (RPL10A-mKate2^lo^GFP^+^) and Ribo-Transferhi (RPL10A-mKate2^hi^GFP^+^) tumor cells were then sorted by FACS and subjected to genomic DNA extraction using the QIAamp DNA Mini Kit (Qiagen). sgRNA cassettes were amplified by two rounds of PCR using primers targeting the GeCKO v2 backbone with Illumina adapters and sample-specific barcodes. Sequencing libraries were gel-purified and sequenced on an Illumina NovaSeq 6000 platform and analyzed using the iScreenAnlys platform provided by ubigene. Raw sequencing reads were mapped to the GeCKO v2 sgRNA reference and analyzed using the MAGeCK algorithm (v0.5.9)^98^. *P*<0.05 was used as the cutoff to define significant hits. Pathway enrichment was performed using gProfiler (https://biit.cs.ut.ee/gprofiler/).

### Public glioma clinical datasets

For comparison of *INTS12* expression between GBM and normal brain tissues, RNA-seq data from TCGA_LGG&GBM and normal brain cortex samples from GTEx (brain, cortex; brain, anterior cingulate cortex (Ba24); and brain, frontal cortex (Ba9)) were retrieved from UCSC Xena (https://xenabrowser.net/datapages/). IDH-wildtype glioma cases were reclassified as GBM according to the 2021 WHO classification^99^, and samples with unknown IDH status were excluded from the original GBM cohort. Pan-cancer analysis of *INTS12* expression was performed using TIMER2.0^100^ (https://compbio.cn/timer2/). For signature and *INTS12* expression analysis in TCGA_LGG&GBM and CGGA datasets, RNA-seq data and clinical information were downloaded from the CGGA website (https://www.cgga.org.cn/). Signature scores were computed using single-sample gene set enrichment analysis (ssGSEA) and converted to Z-scores using the formula: Z-score = (expression value - mean) / standard deviation, where the mean and standard deviation were calculated across all samples for each gene or gene set. The Ribo-Transferʰⁱ signature was derived from top 50 upregulated genes in the Ribo-Transferʰⁱ cells identified from mouse single-cell RNA-seq data using the FindMarkers function (Seurat package (v5.5.0), ranked by log_2_ fold change). Human orthologs were obtained using the babelgene (v22.9) R package, and non-orthologous genes were excluded. The list of signature genes is provided in Table S4.

## QUANTIFICATION AND STATISTICAL ANALYSIS

All statistical analyses are described in figure legends. Statistical tests, including two-tailed unpaired t test, two-tailed paired t test, one-way ANOVA, two-way ANOVA, two-tailed Pearson correlation, log-rank test, two-tailed Wilcoxon rank-sum test and Kruskal-Wallis test were performed using R (v4.5.3) or GraphPad Prism (v9.5.0) software. Unless otherwise indicated, data in the figures are presented as mean ± SD. For all statistical tests, *P* < 0.05 was taken to indicate statistical significance unless otherwise stated.

---

**Figure S1.**
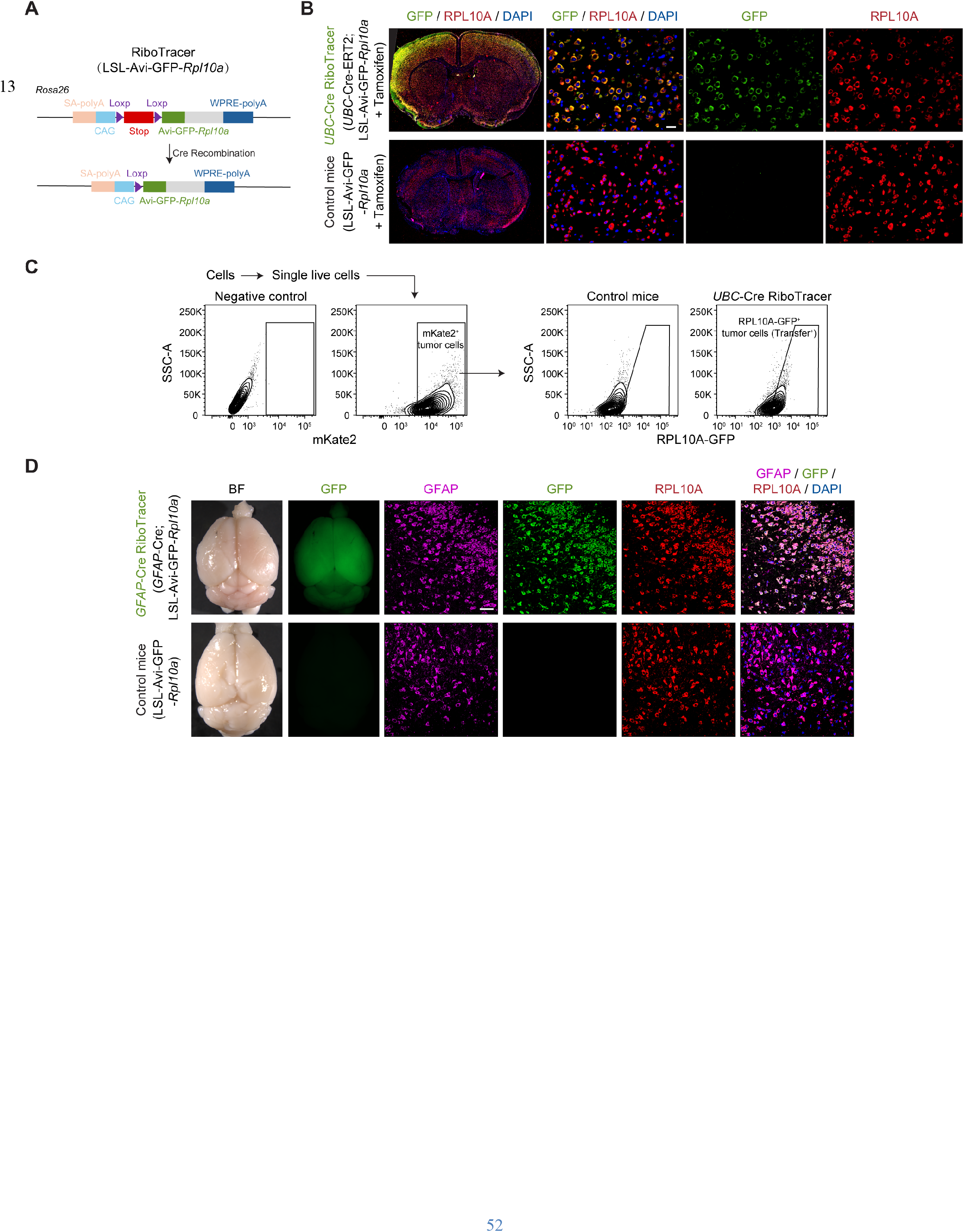
Characterization of the RiboTracer reporter mouse, related to Figure 1. (A) Schematics of the RiboTracer reporter mouse. (B) Representative images of GFP and RPL10A staining in *UBC*-Cre; RiboTracer mice and controls. Scale bar, 20 μm. (C) Gating strategy used to identify ribosome transfer (RPL10A-GFP signal in mKate2^+^ tumor cells) in tumor-bearing *UBC*-Cre; RiboTracer reporter mouse. (D) Representative images of GFP, GFAP and RPL10A staining in *GFAP*-Cre; RiboTracer mice and controls. Scale bar, 50 μm.

**Figure S2.**
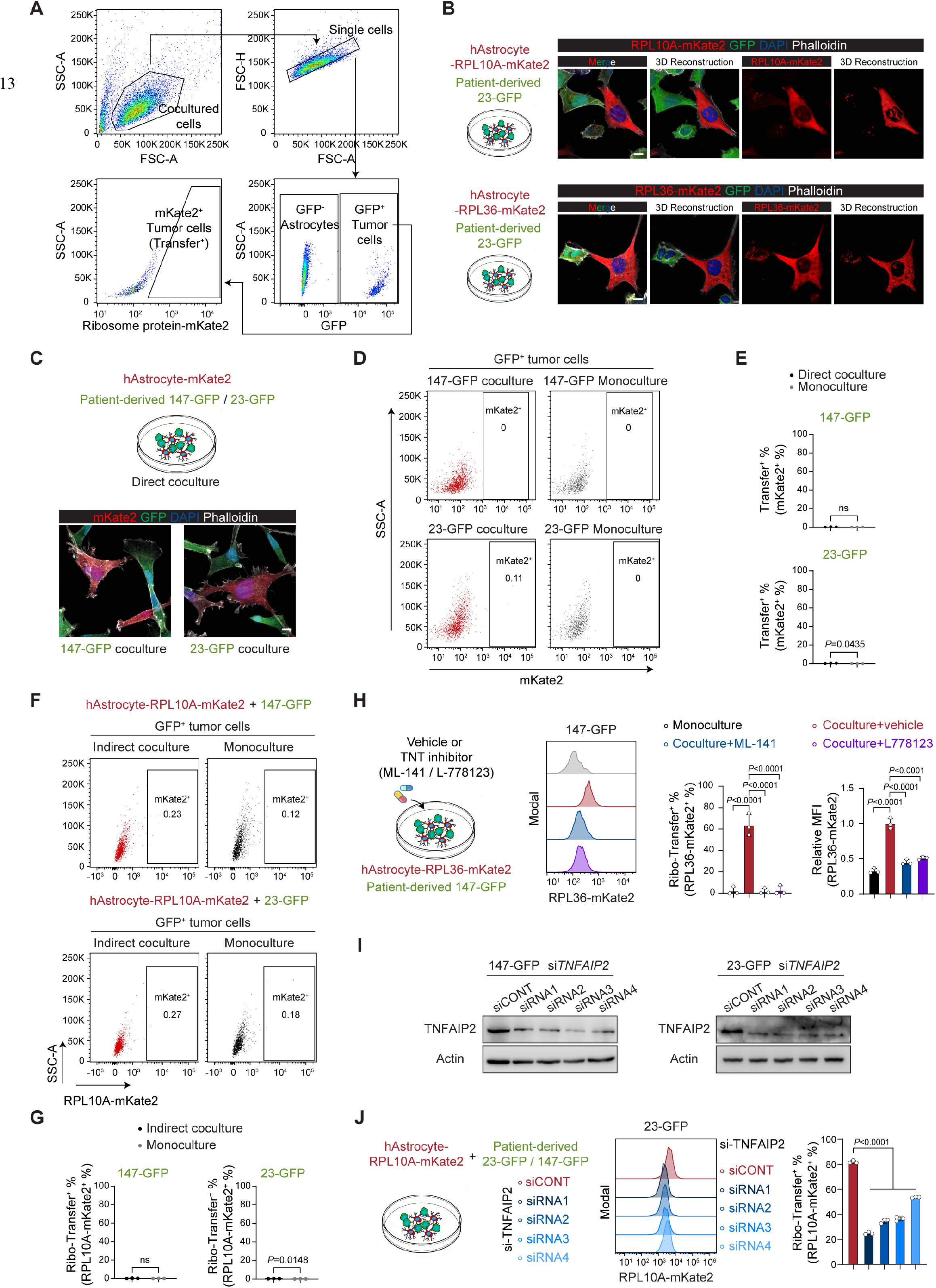
Direct ribosome transfer via tunneling nanotube-like structures, related to Figure 2. (A) Gating strategy used to identify ribosome transfer (ribosome protein-mKate2 signal in GFP^+^ tumor cells) in the coculture system. (B) Schematics and representative images of RPL10A-mKate2 or RPL36-mKate2 in the indicated coculture system. GFP labels tumor cells; phalloidin labels F-actin (TNT marker). Scale bar, 10 μm. (C) Schematics and representative images of fluorescent proteins transfer in direct coculture of human astrocytes expressing mKate2 alone and GFP-labeled patient-derived tumor cells. Scale bar, 10 μm. (**D** and **E**) Representative flow plots and quantification of mKate2 in GFP^+^ tumor cells in the direct coculture from (C). (**F** and **G**) Representative flow plots and quantification of RPL10A-mKate2 in GFP^+^ tumor cells in indirect coculture separated by a transwell chamber. (H) Schematics, representative flow plots, and quantification of RPL36-mKate2 in direct coculture treated with 10 μM ML-141, 10 μM L-778123, or vehicle (n=3 per group). (I) Immunoblots showing TNFAIP2 protein levels in patient-derived tumor cells treated with siRNAs targeting *TNFAIP2* or control siRNA. (J) Schematics, representative flow plots, and quantification of RPL10A-mKate2 in direct coculture of human astrocytes expressing RPL10A-mKate2 and 23-GFP treated with the indicated siRNAs targeting *TNFAIP2* (n=3 per group). Data are represented as the mean ± SD in (E), (G), (H) and (J). Individual data points represent independent experiments. Two-tailed unpaired t test for (E) and (G). One-way ANOVA followed by multiple comparisons for (H) and (J). ns, not significant.

**Figure S3.**
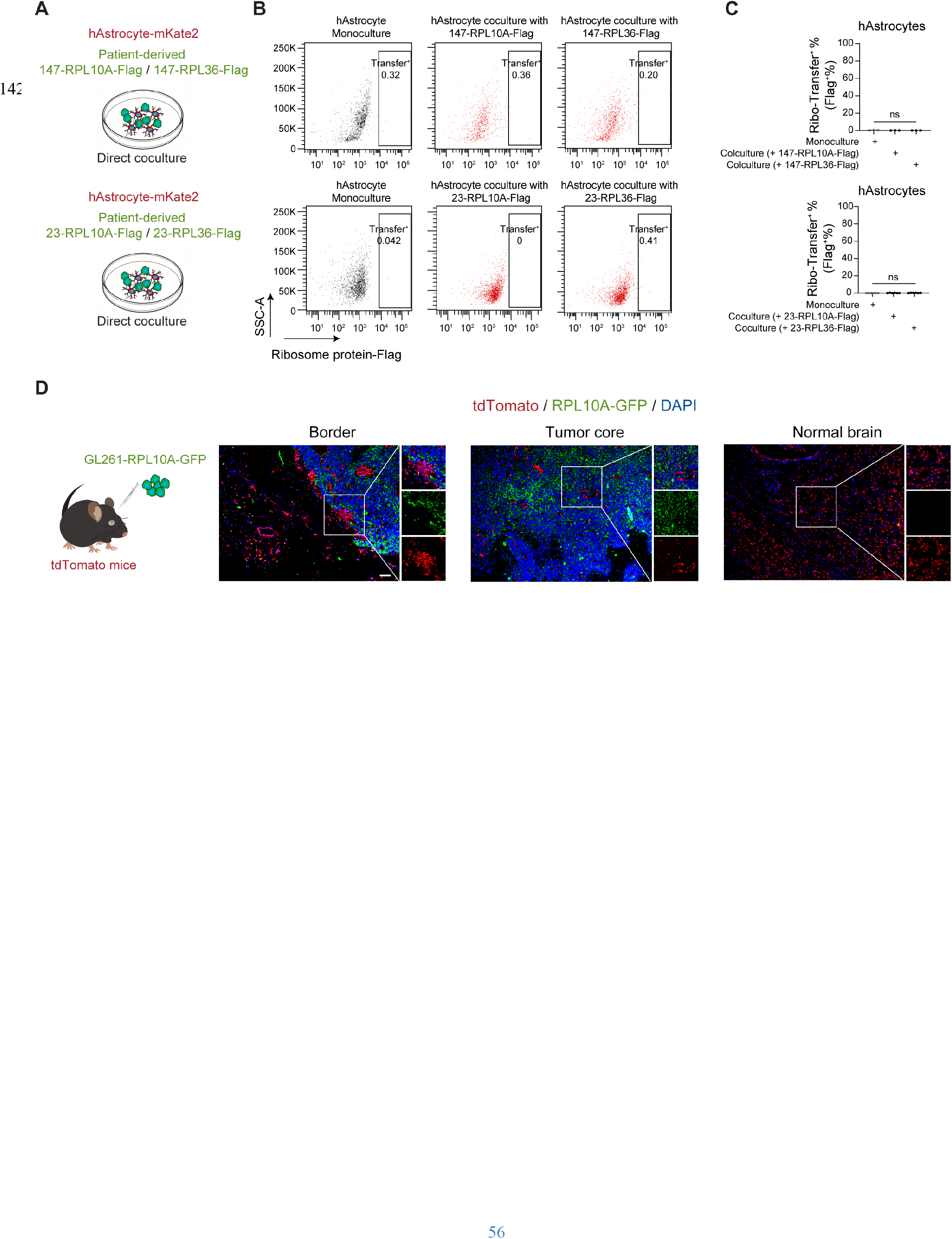
Ribosomes are transferred unidirectionally from astrocytes to GBM cells, related to Figure 2. (**A-C**) Schematics of reciprocal labeling, representative flow plots, and quantification of RPL10A- Flag or RPL36-Flag in mKate2-labeled human astrocytes cultured alone or directly cocultured with patient-derived #23 or #147 GBM cells expressing RPL10A-Flag or RPL36-Flag (n=3 independent experiments per group). Data are represented as the mean ± SD. Statistical significance was determined by one-way ANOVA followed by multiple comparisons. ns, not significant. (D) Schematics of reciprocal labeling *in vivo* and representative images of RPL10A-GFP in brain slices from GL261-bearing tdTomato reporter mice. tdTomato labels non-malignant cells. Scale bar, 50 μm.

**Figure S4.**
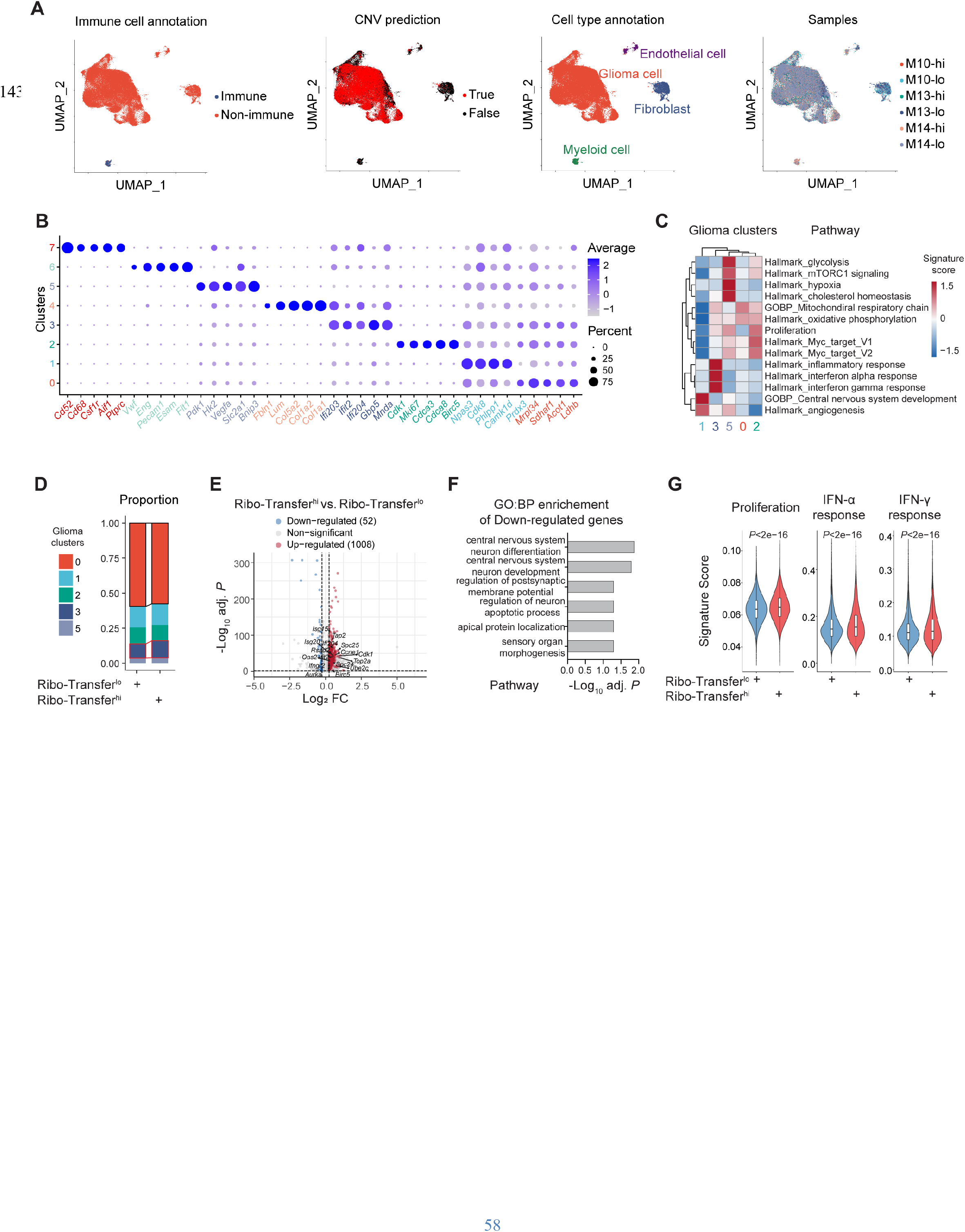
scRNA-seq of Ribo-Transfer^lo^ and Ribo-Transfer^hi^ GL261 cells, related to Figure 4. (A) UMAP plots showing the immune cell annotation, CNV prediction, cell type annotation and sample distribution in scRNA-seq of mouse GBM samples. (B) Dot plot of marker gene expression across clusters. (C) Heatmap showing indicated pathway scores across glioma clusters. (D) Stacked bar plot showing the proportion of glioma clusters in Ribo-Transfer^hi^ vs. Ribo- Transfer^lo^ GL261 cells. (E) Volcano plot showing differentially expressed genes in Ribo-Transfer^hi^ vs. Ribo-Transfer^lo^ GL261 cells from scRNA-seq. Selected interferon-stimulated genes (ISGs) and cell cycle-related genes are highlighted. (F) GO:BP enrichment of downregulated genes in Ribo-Transfer^hi^ vs. Ribo-Transfer^lo^ GL261 cells from scRNA-seq. Statistical significance was determined by overrepresentation test corrected by FDR. (G) Violin plots showing indicated signature scores in Ribo-Transfer^lo^ (n=28,990 cells) and Ribo- Transfer^hi^ (n=19,450 cells) GL261 cells from scRNA-seq. Violin plots show the overall distribution of data points; boxes indicate median, upper, and lower quartiles. Statistical significance was determined by two-tailed Wilcoxon rank-sum test.

**Figure S5.**
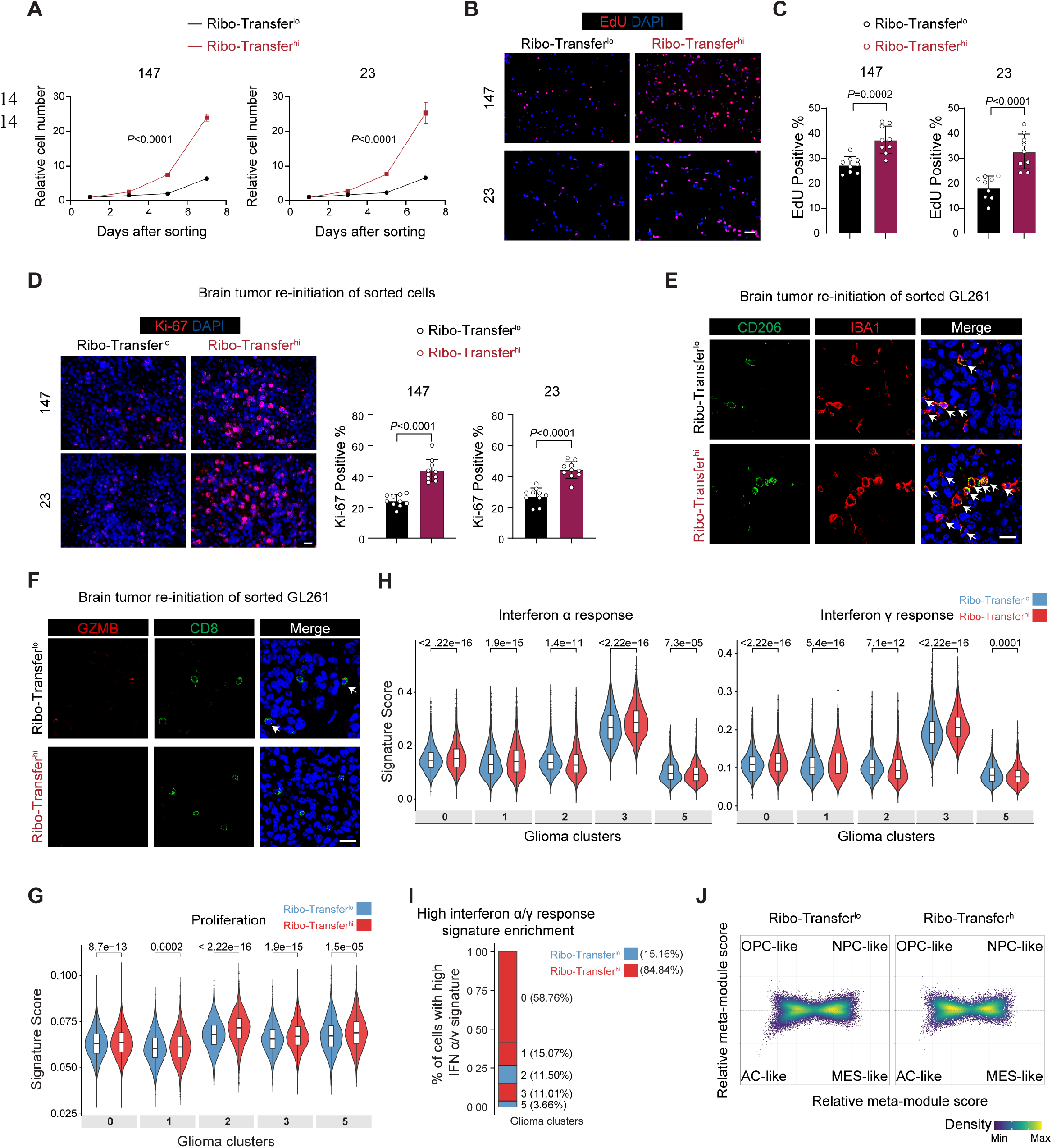
Ribosome transfer enhances cancer cell proliferation and triggers an IFN response, related to Figure 4. (A) Normalized cell numbers of Ribo-Transferˡᵒ and Ribo-Transferʰⁱ patient-derived #23 and #147 GBM cells at the indicated time points post-sorting (n=3 independent experiments). Data are normalized to day 1. (**B** and **C**) Representative images and quantification (n=9 randomly selected fields from three independent experiments) of EdU incorporation in Ribo-Transferˡᵒ and Ribo-Transferʰⁱ #23 and #147 GBM cells. Scale bar, 50 μm. (D) Representative and quantification of Ki-67^+^ percentage (n=10 randomly selected fields from at least three mice) in brain sections from tumor-bearing mice initiated by sorted Ribo-Transfer^lo^ or Ribo-Transfer^hi^ #23 and #147 GBM cells. Scale bar, 20 μm. (**E** and **F**) Representative staining of suppressive myeloid cells (IBA1^+^CD206^+^) and effector T cells (CD8^+^GZMB^+^) in brain sections from tumor-bearing mice implanted with sorted Ribo- Transfer^lo^ or Ribo-Transfer^hi^ GL261 cells. Quantification is shown in Fig. 4I. Arrows indicate double positive cells. Scale bar, 20 μm. (**G** and **H**) Violin plots showing indicated signature scores in Ribo-Transfer^lo^ (n=28,990 cells) and Ribo-Transfer^hi^ (n=19,450 cells) GL261 cells across glioma clusters from scRNA-seq. Violin plots show the overall distribution of data points; boxes indicate median, upper, and lower quartiles. (I) Stacked bar plot showing the proportion of glioma clusters with high IFN α/γ response (Ribo- Transfer^hi^ vs. Ribo-Transfer^lo^). (J) Four-quadrant plot showing the distribution of OPC-like, NPC-like, AC-like and MES-like cell states in Ribo-Transfer^lo^ and Ribo-Transfer^hi^ GL261 cells from scRNA-seq. Data are represented as the mean ± SD in (A), (C) and (D). Two-way ANOVA followed by multiple comparison for (A). Two-tailed unpaired t test for (C) and (D). Two-tailed Wilcoxon rank-sum test for (H) and (G).

**Figure S6.**
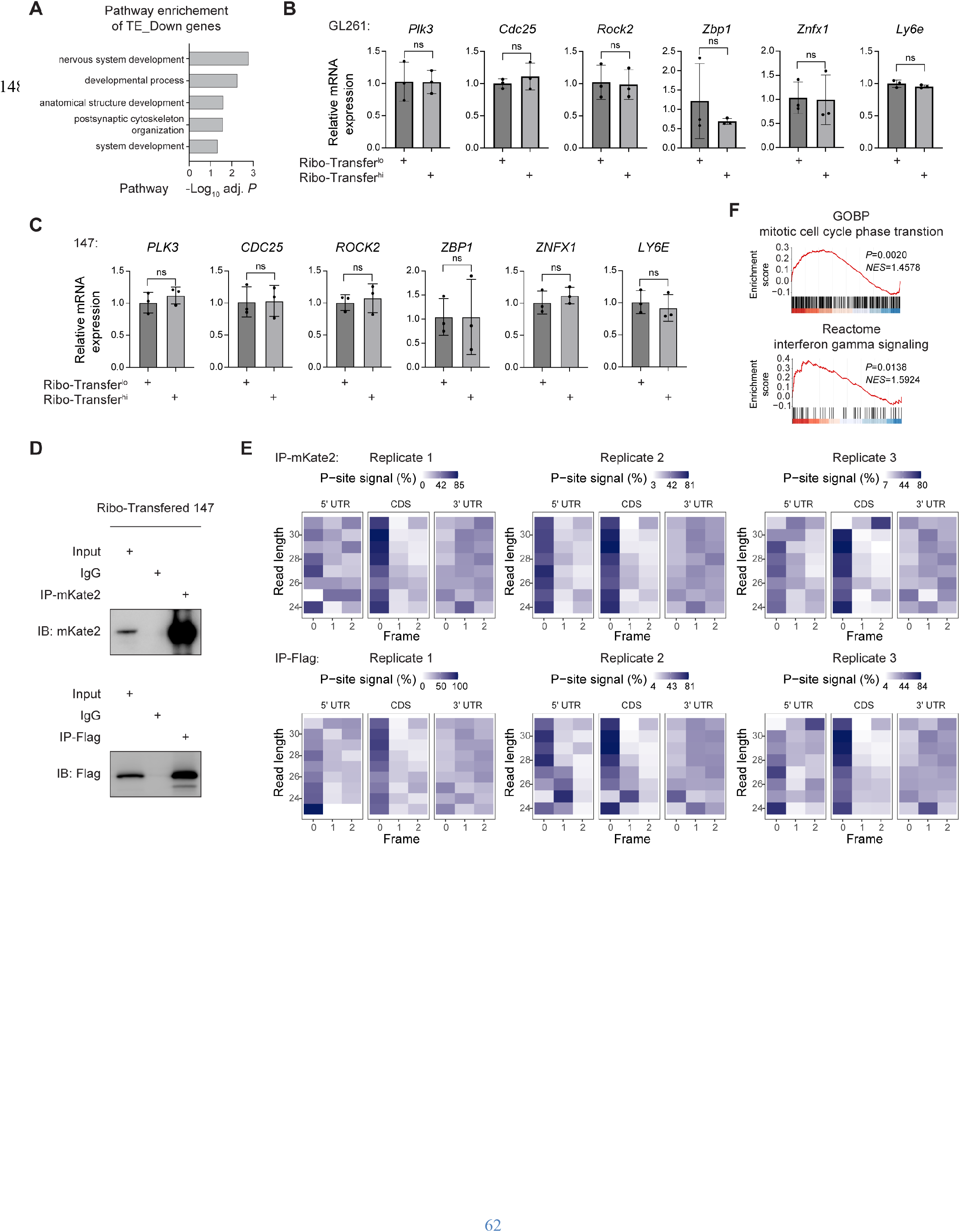
Astrocyte-derived ribosomes are functionally engaged in protein translation, related to Figure 4. (A) GO:BP pathway enrichments of genes with downregulated translation efficiency (TE) in Ribo- Transfer^hi^ vs. Ribo-Transfer^lo^ GL261 cells. (**B** and **C**) qPCR analysis of indicated genes in Ribo-Transfer^lo^ and Ribo-Transfer^hi^ GL261 cells isolated from orthotopic tumors and #147 cells isolated from coculture (n=3 independent experiments). (**D**) Immunoblots showing mKate2 expression in IP-mKate2 and Flag expression in IP-Flag. (**E**) Heatmap showing percentage of P-site signals on three-nucleotide (0, 1, 2) codon frames in 5’UTR, CDS and 3’UTR in Ribosome-seq of IP-mKate2 and IP-Flag. (**F**) Representative GSEA plots showing significant pathways enriched in IP-mKate2. Data are represented as the mean ± SD in (B) and (C). Overrepresentation test corrected by FDR for (A). Two-tailed unpaired t test for (B) and (C). Weighted Kolmogorov–Smirnov statistic test for (E). ns, not significant.

**Figure S7.**
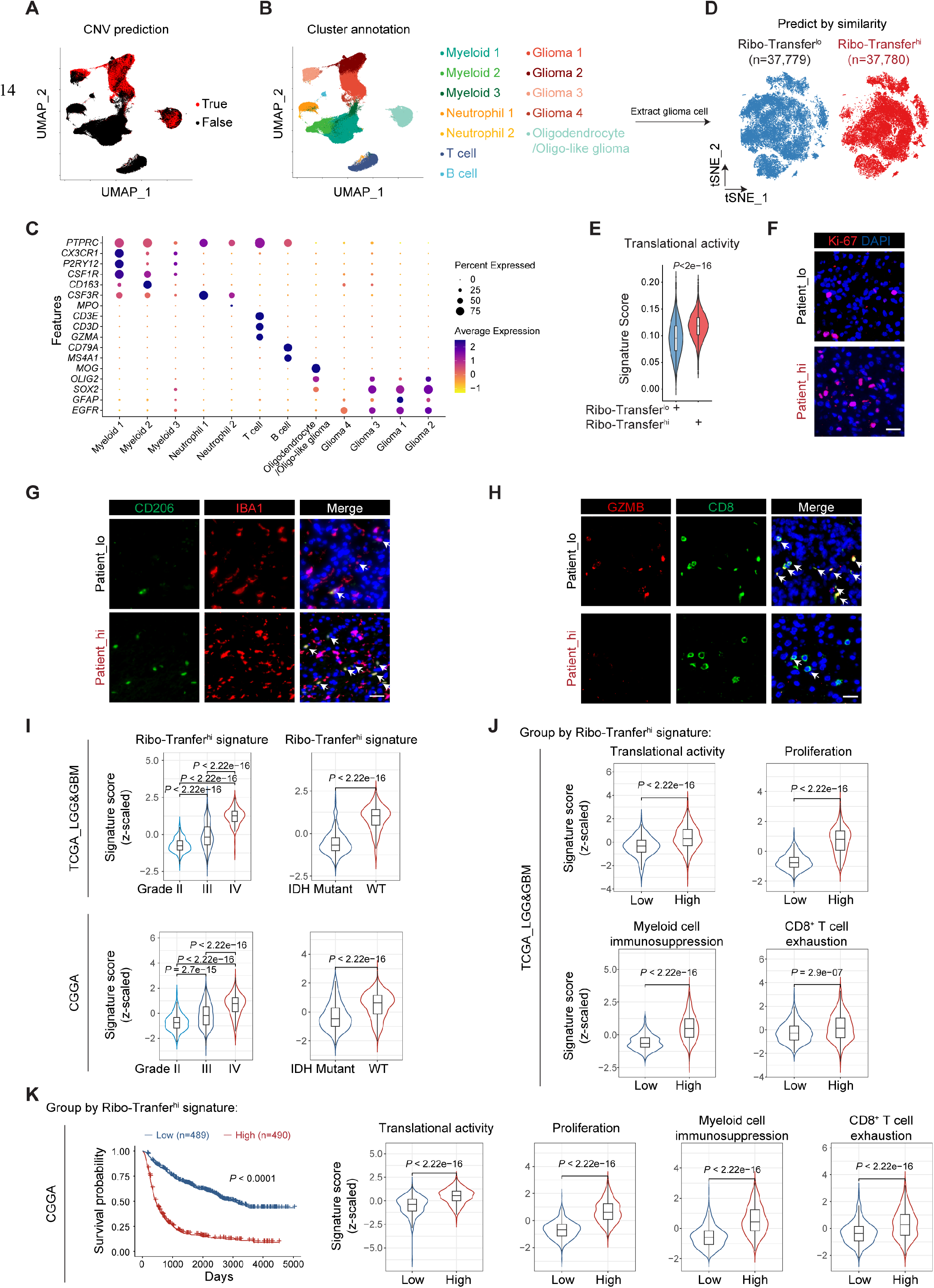
Clinical relevance of ribosome transfer, related to Figure 5. (**A** and **B**) UMAP plots showing the CNV prediction, cluster annotation in scRNA-seq of 20 human GBM samples. (C) Dot plot of marker gene expression across clusters. (D) *t*-SNE plots showing the distribution of predicted Ribo-Transfer^lo^ and Ribo-Transfer^hi^ human GBM cells. (E) Violin plots showing translational activity signature scores in Ribo-Transfer^lo^ (n=37,779 cells) and Ribo-Transfer^hi^ (n=37,780 cells) GBM cells from scRNA-seq. (**F-H**) Representative staining of Ki-67, suppressive myeloid cells (IBA1^+^CD206^+^) and effector T cells (CD8^+^GZMB^+^) in brain sections from human GBM patients. Quantification is shown in Fig. 5F. Arrows indicate double positive cells. Scale bar, 20 μm. (I) Violin plots showing Ribo-Transfer^hi^ signature score across tumor grade and IDH status in TCGA_LGG&GBM (Grade II, n=216; Grade III, n=241; Grade IV, n=152; IDH Mutant: n=428; IDH WT: n=234) and CGGA (Grade II, n=291; Grade III, n=334; Grade IV, n=388; IDH Mutant: n=531; IDH WT: n=435) datasets. (J) Violin plots showing indicated signature score in patients stratified by low (n=304) and high (n=305) Ribo-Transfer^hi^ signature in TCGA_LGG&GBM dataset. Cutoff, median ssGSEA signature score. (K) Kaplan–Meier survival curves and violin plots showing indicated signature score in patients stratified by low (n=489) vs. high (n=490) Ribo-Transfer^hi^ signature in the CGGA dataset. Cutoff, median ssGSEA signature score. In (E), (I) to (K), violin plots show the overall distribution of data points; boxes indicate median, upper, and lower quartiles. The number of n represents patients unless otherwise specified. Two-tailed Wilcoxon rank-sum test for (E), (I) to (K). Kruskal-Wallis test followed by multiple comparisons for (I). Log-rank test for (K).

**Figure S8.**
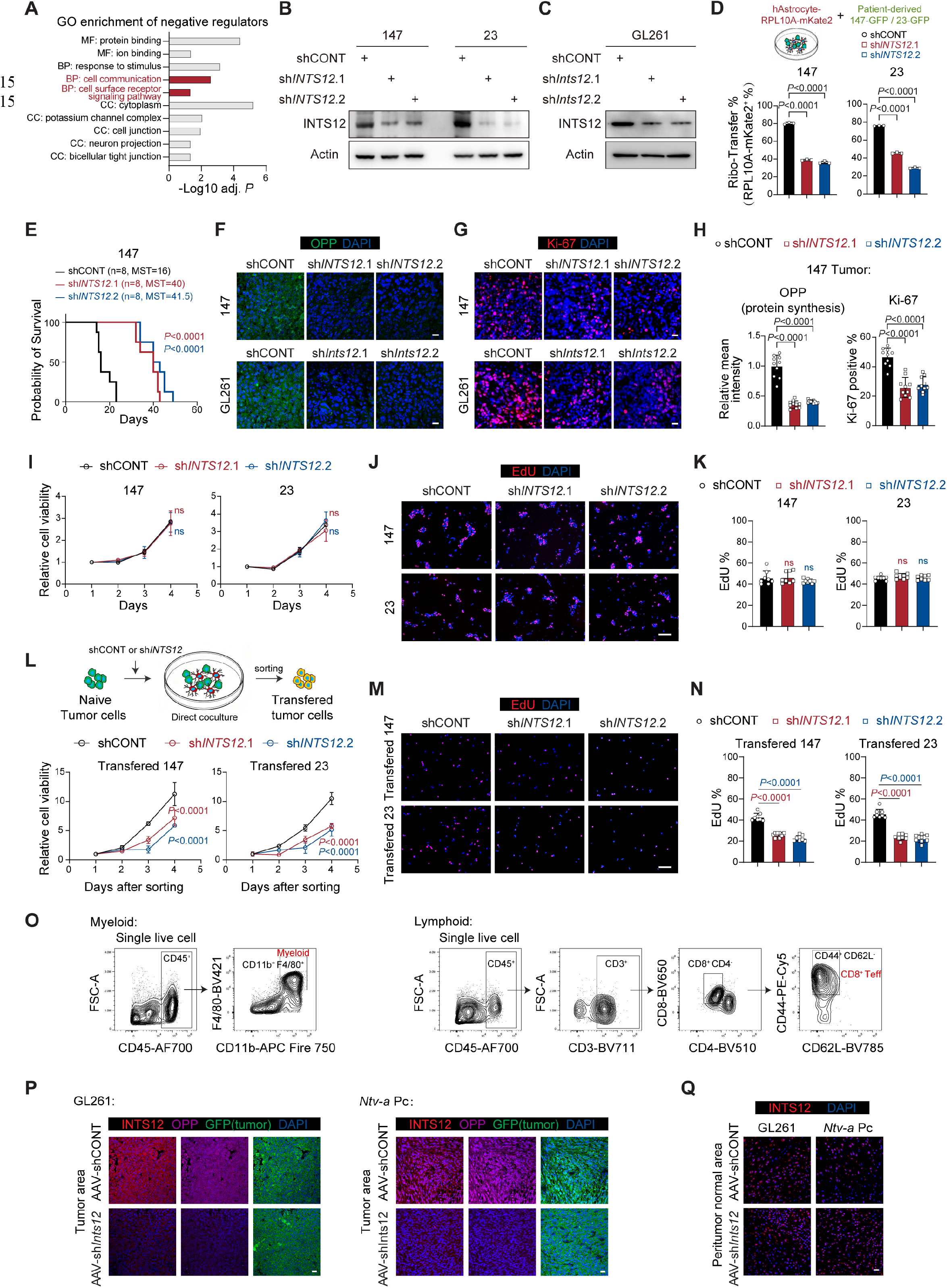
INTS12 promotes proliferation via non-cell-autonomous ribosome transfer, related to Figure 5. (**A**) GO enrichment of significant hits in negative selection (Ribo-Transfer^hi^ vs. Ribo-Transfer^lo^) of the CRISPR screen. (**B** and **C**) Immunoblots showing INTS12 protein levels in #23 and #147 GBM cells with INTS12 knockdown and GL261 cells with Inst12 knockdown. (D) Schematics and quantification of RPL10A-mKate2 in direct coculture of human astrocyte expressing RPL10A-mKate2 and 147-GFP or 23-GFP with or without *INTS12* knockdown (n=3 independent experiments). (E) Kaplan–Meier survival curves of nude mice implanted with #147 GBM cells with or without *INTS12* knockdown (n=8 mice per group). MST, median survival time. (**F** and **G**) Representative staining of OPP incorporation and Ki-67 in #147 GBM cells with or without *INTS12* knockdown and GL261 cells with or without *Inst12* knockdown. Scale bar, 20 μm. (H) Quantification of OPP incorporation and Ki-67^+^ percentage (n=10 randomly selected fields from at least three mice) in #147 GBM cells with or without *INTS12* knockdown. (I) Relative cell viability of #23 and #147 GBM cells with or without *INTS12* knockdown at the indicated time points (n=3 independent experiments). Data are normalized to day 1. (**J** and **K**) Representative images and quantification (n=8 randomly selected fields from at least three independent experiments) of EdU incorporation in #23 and #147 GBM cells with or without *INTS12* knockdown. Scale bar, 50 μm. (L) Relative cell viability of ribosome tranferred #23 and #147 GBM cells with or without *INTS12* knockdown at the indicated time points (n=3 independent experiments). Schematics showing the sorting strategy. Data are normalized to day 1. (**M** and **N**) Representative images and quantification (n=8 randomly selected fields from at least three independent experiments) of EdU incorporation in ribosome tranferred #23 and #147 GBM cells with or without *INTS12* knockdown at the indicated time points. Scale bar, 50 μm. (**O**) Gating strategy to identify myeloid cells and CD8^+^ effector T cells in flow cytometry. (**P** and **Q**) Representative staining of INTS12 and OPP incorporation in brain sections from GL261- or *Ntv-a* Pc-bearing mice treated with AAV-shCONT or AAV-sh*Ints12.* Tumor cells are labelled with GFP to identify tumor areas. Scale bar, 20 μm. Data are represented as the mean ± SD in (D), (H), (I), (K), (L) and (N). Overrepresentation test corrected by FDR for (A). One-way ANOVA followed by multiple comparisons for (D), (H), (K) and (N). Log-rank test for (E). Two-way ANOVA followed by multiple comparisons for (I) and (L). ns, not significant.

**Figure S9.**
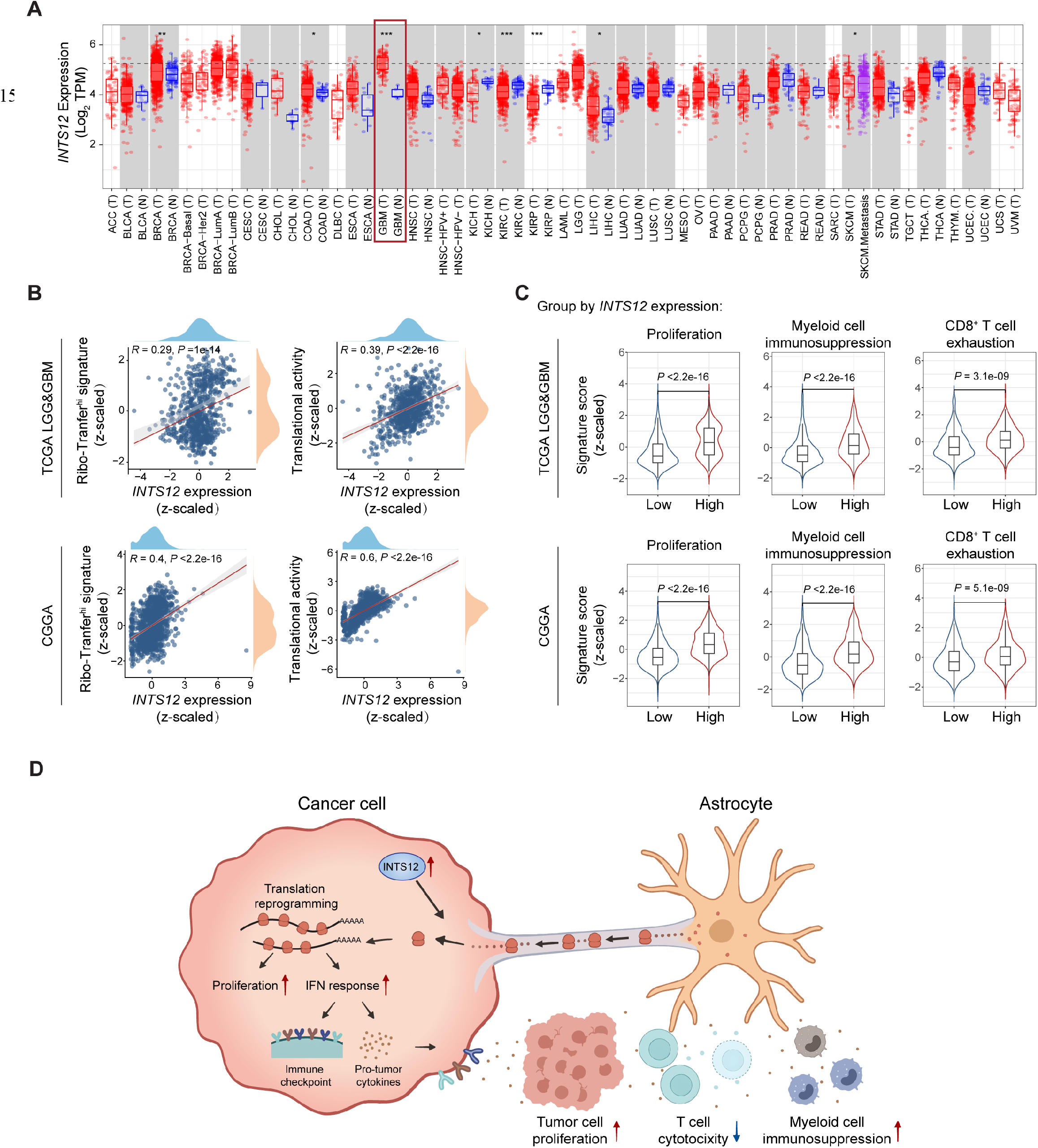
Clinical relevance of INTS12, related to Figure 5. (A) Box plot showing *INTS12* expression across TCGA pan-cancer cohorts. The statistical significance computed by differential analysis (edgeR) on RNA-Seq raw counts is annotated by the number of stars (*: *P*-value < 0.05; **: *P* -value <0.01; ***: *P* -value <0.001). (B) Pearson correlation of INTS12 and Ribo-Transfer^hi^ or translation signature in TCGA_LGG&GBM (n=702) and CGGA (n=1,008) datasets. Statistical significance was determined by two-tailed Pearson correlation. (C) Violin plots showing *INTS12* expression in patients stratified by low and high Ribo-Transfer^hi^ signature in TCGA_LGG&GBM (low: n=304; high: n=305) and CGGA (low: n=489; high: n=490) dataset. Cutoff, median ssGSEA signature score. Violin plots show the overall distribution of data points; boxes indicate median, upper, and lower quartiles. Statistical significance was determined by two-tailed Wilcoxon rank-sum test. (D) Graphic abstract of this study.

**Table S1.** Patient information for scRNA-seq samples.

| Sample | Gender | Age | WHO grade | Primary or recurrent | IDH status |
| --- | --- | --- | --- | --- | --- |
| GBM01 | male | 41 | 4 | primary | wildtype |
| GBM02 | male | 65 | 4 | primary | wildtype |
| GBM03 | male | 56 | 4 | primary | wildtype |
| GBM04 | male | 36 | 4 | primary | wildtype |
| GBM05 | male | 69 | 4 | primary | wildtype |
| GBM06 | male | 59 | 4 | primary | wildtype |
| GBM07 | male | 43 | 4 | primary | wildtype |
| GBM08 | female | 52 | 4 | primary | wildtype |
| GBM09 | male | 71 | 4 | primary | wildtype |
| GBM10 | male | 39 | 4 | primary | wildtype |
| GBM11 | male | 77 | 4 | primary | wildtype |
| GBM12 | female | 71 | 4 | primary | wildtype |
| GBM13 | male | 53 | 4 | primary | wildtype |
| GBM14 | male | 65 | 4 | primary | wildtype |
| GBM15 | female | 47 | 4 | primary | wildtype |
| GBM16 | male | 60 | 4 | primary | wildtype |
| GBM17 | female | 66 | 4 | primary | wildtype |
| GBM18 | male | 57 | 4 | primary | wildtype |
| GBM19 | female | 64 | 4 | primary | wildtype |
| GBM20 | male | 73 | 4 | primary | wildtype |

**Table S2.**
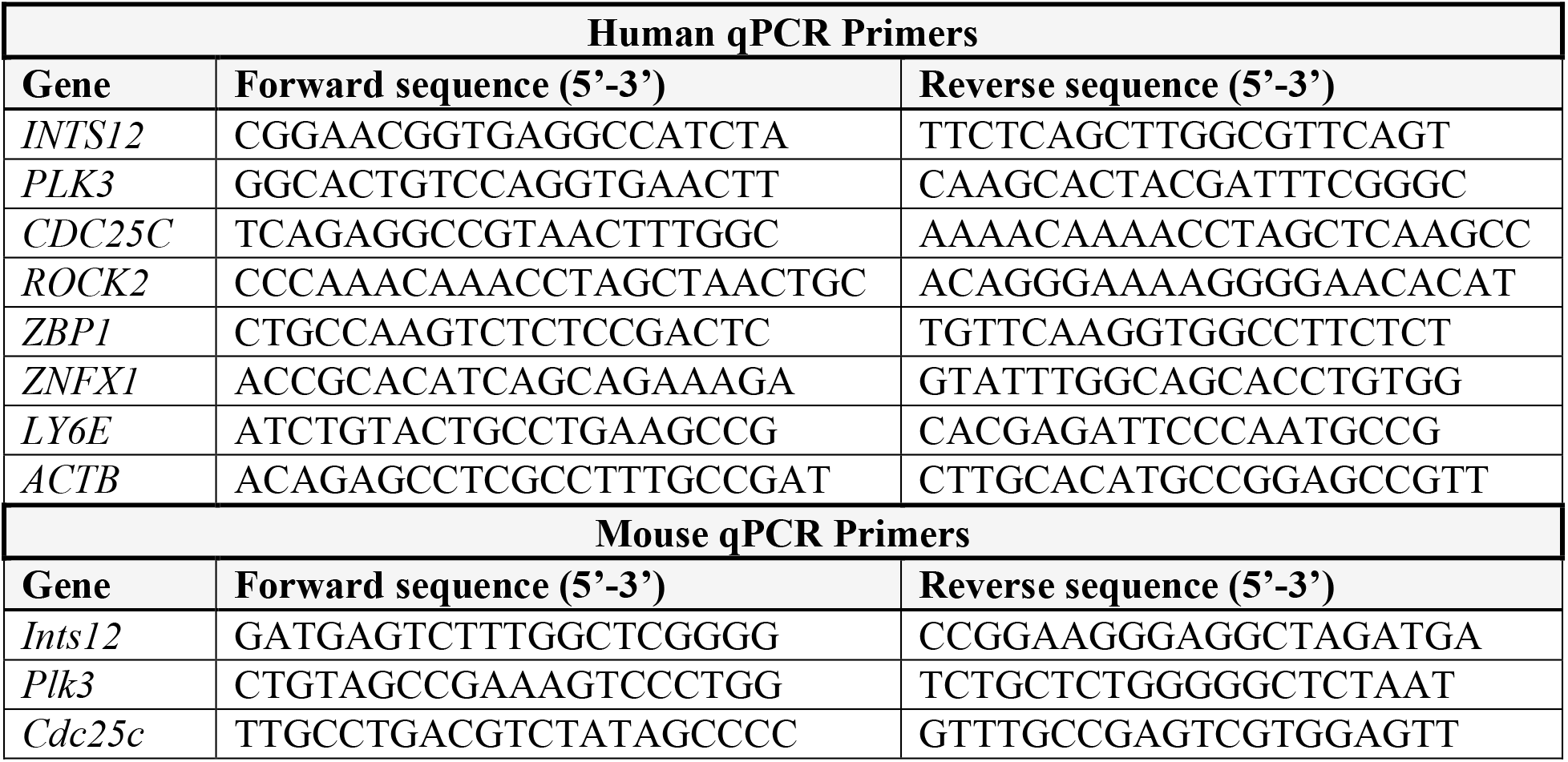
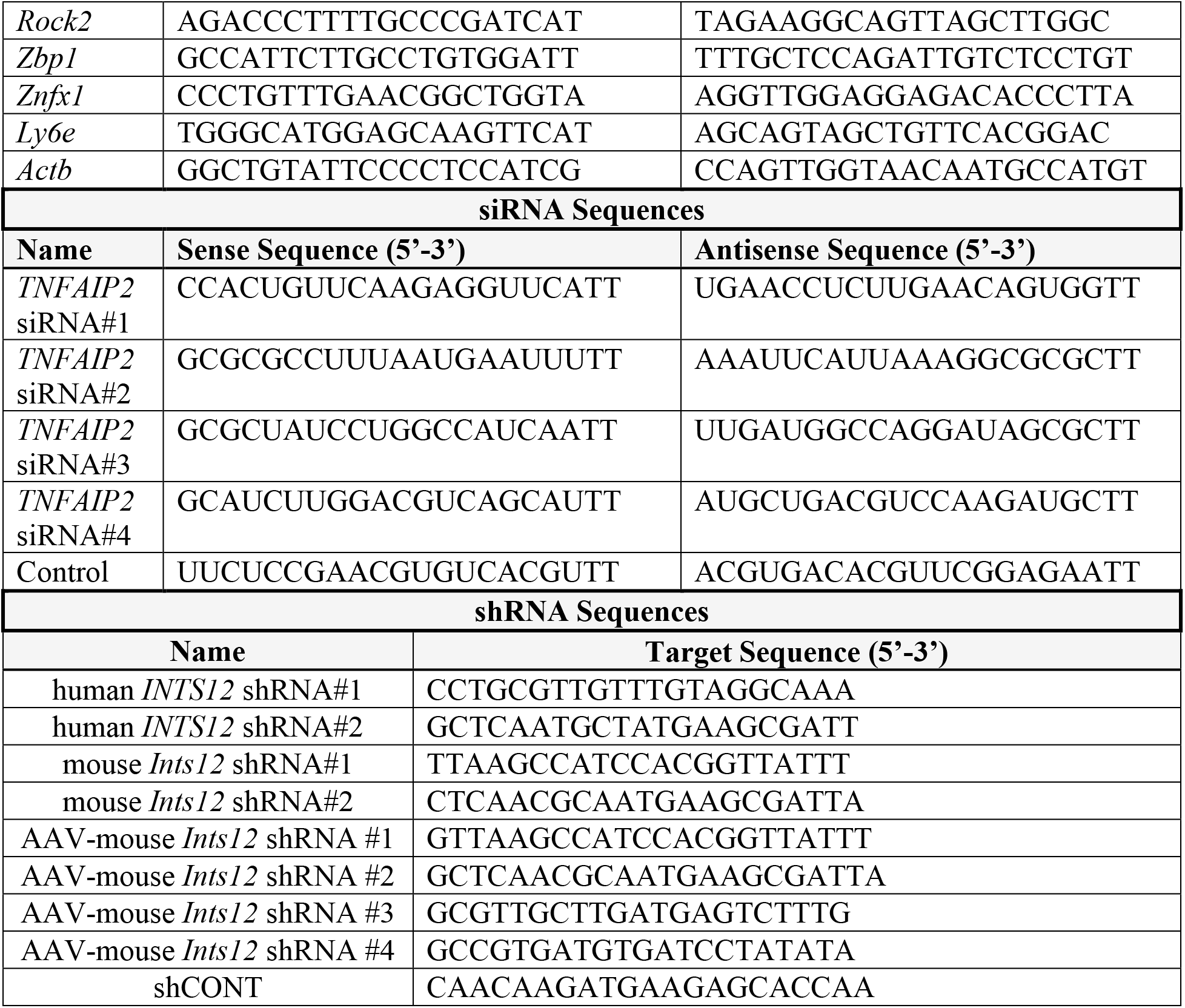
Primers and siRNA/shRNA sequences used in this study.

**Table S3.**
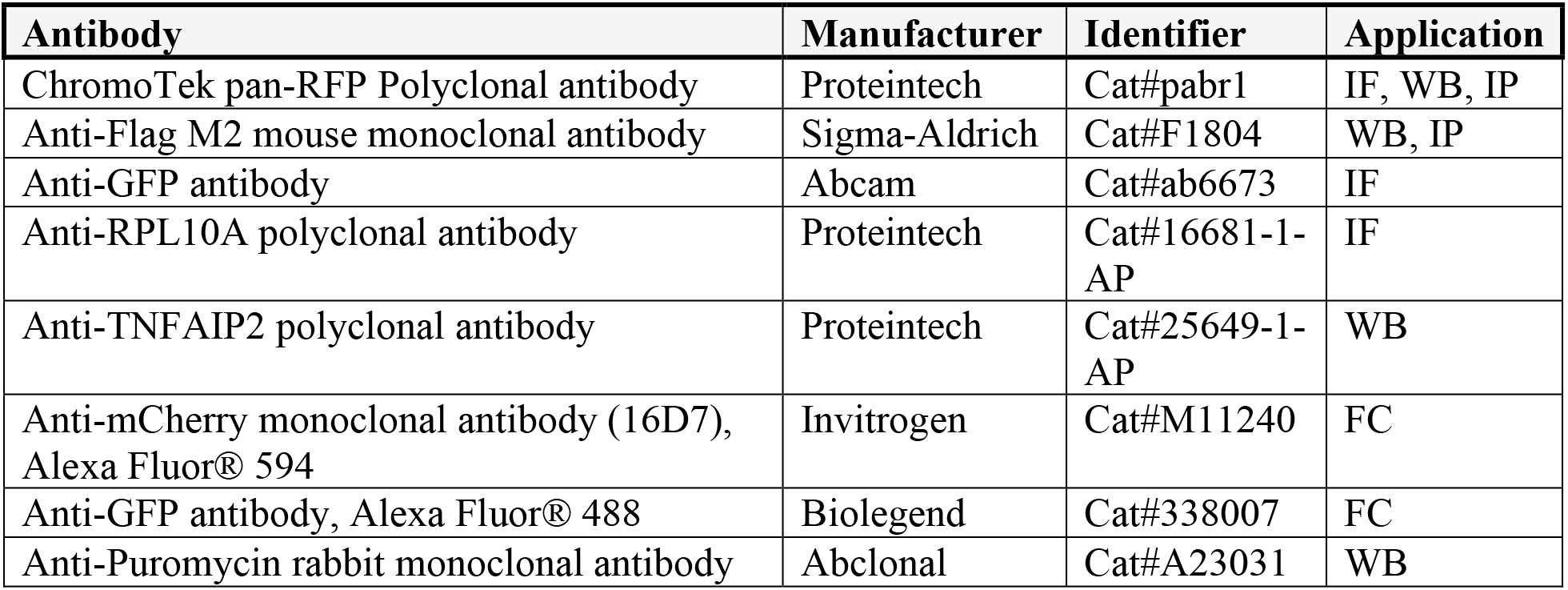
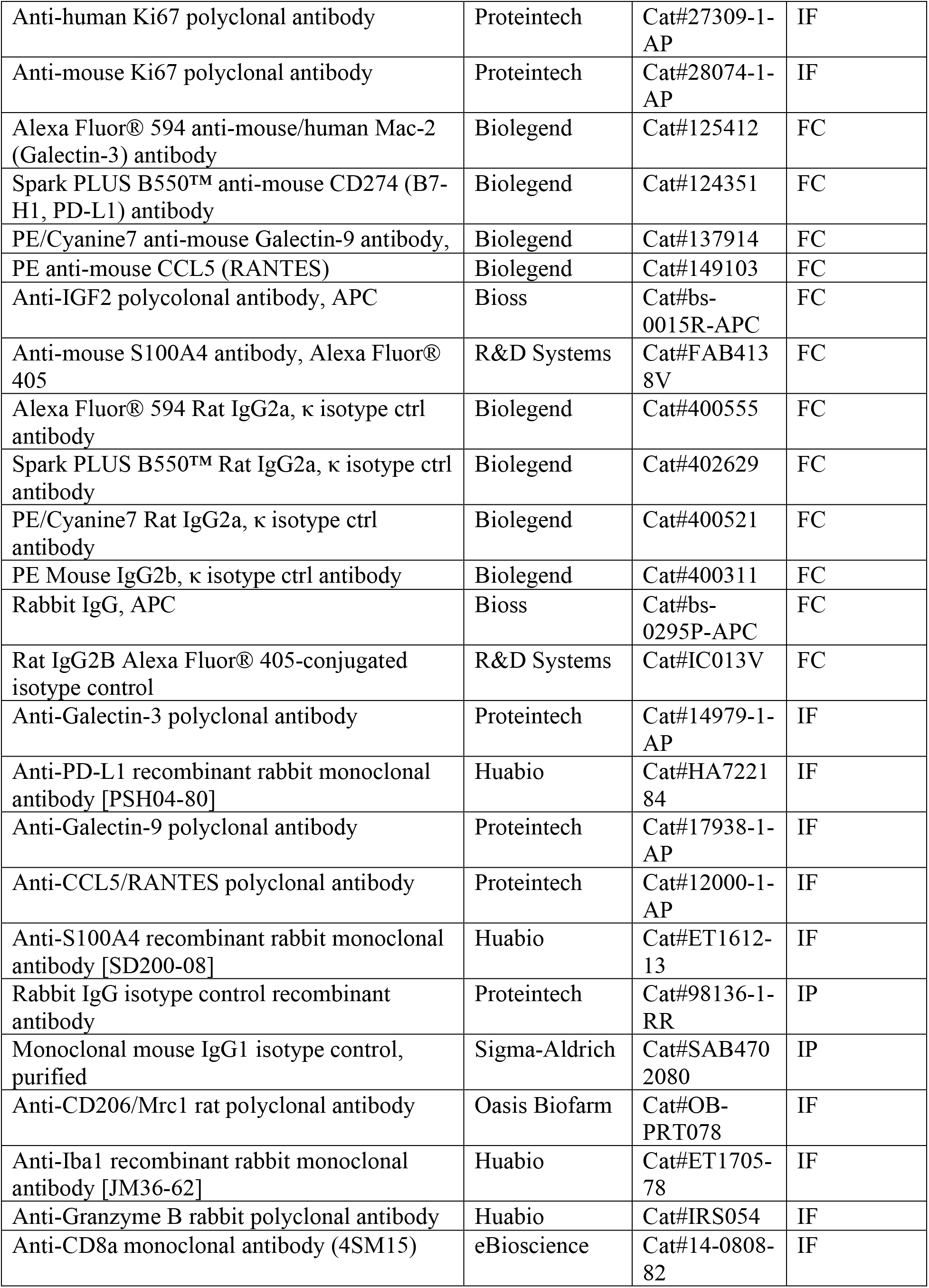
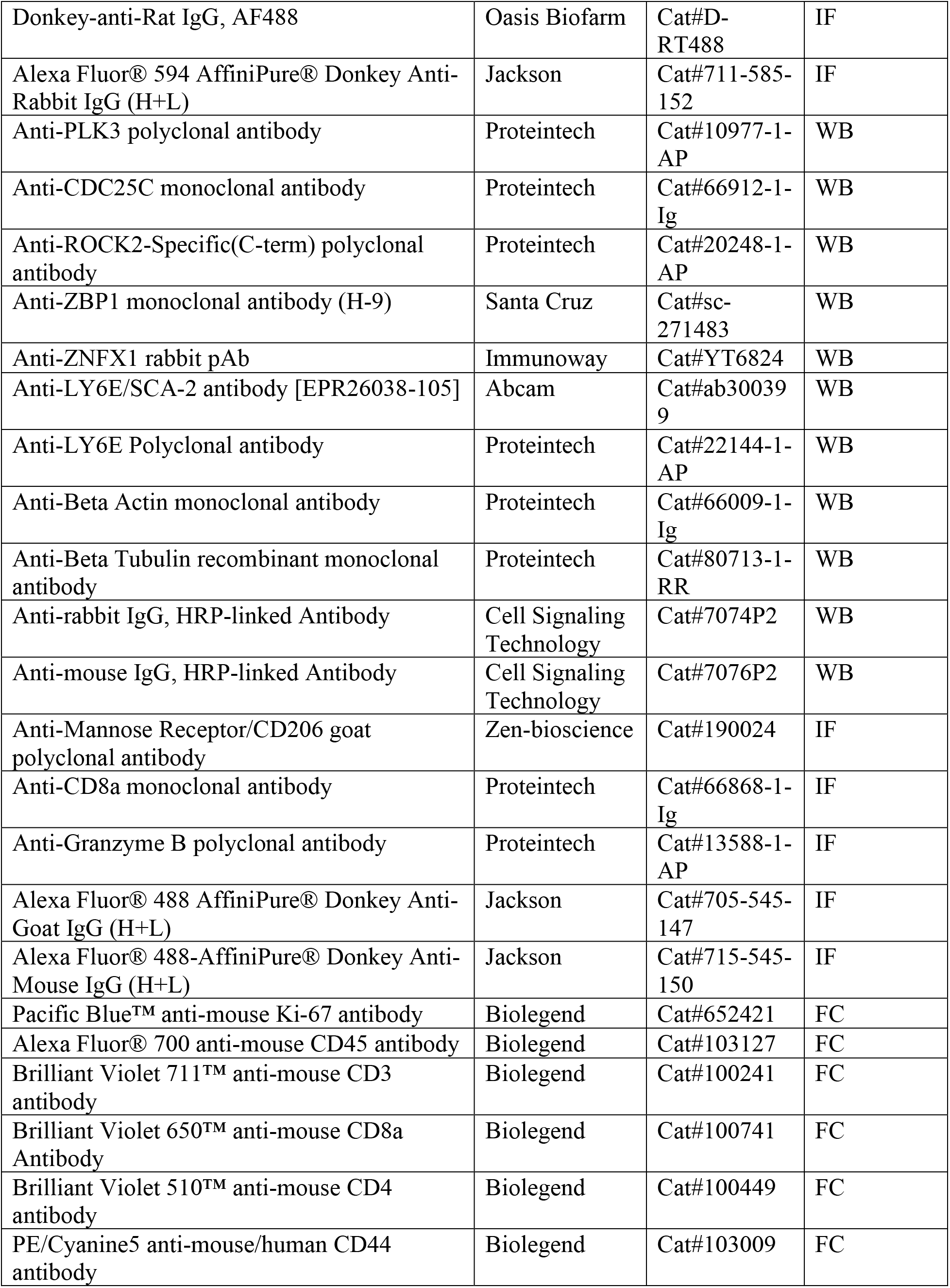
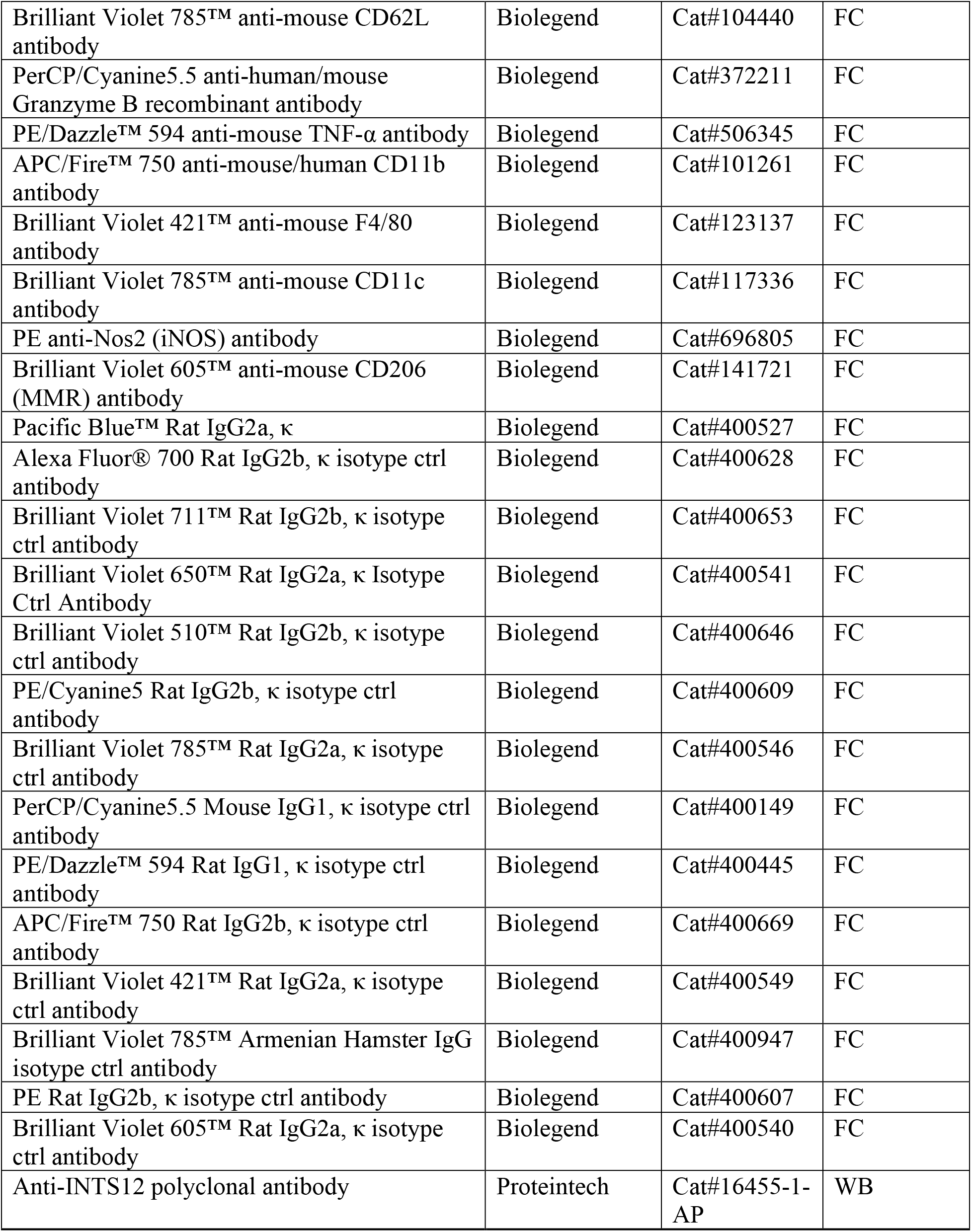
Antibodies used in this study.

**Table S4.**
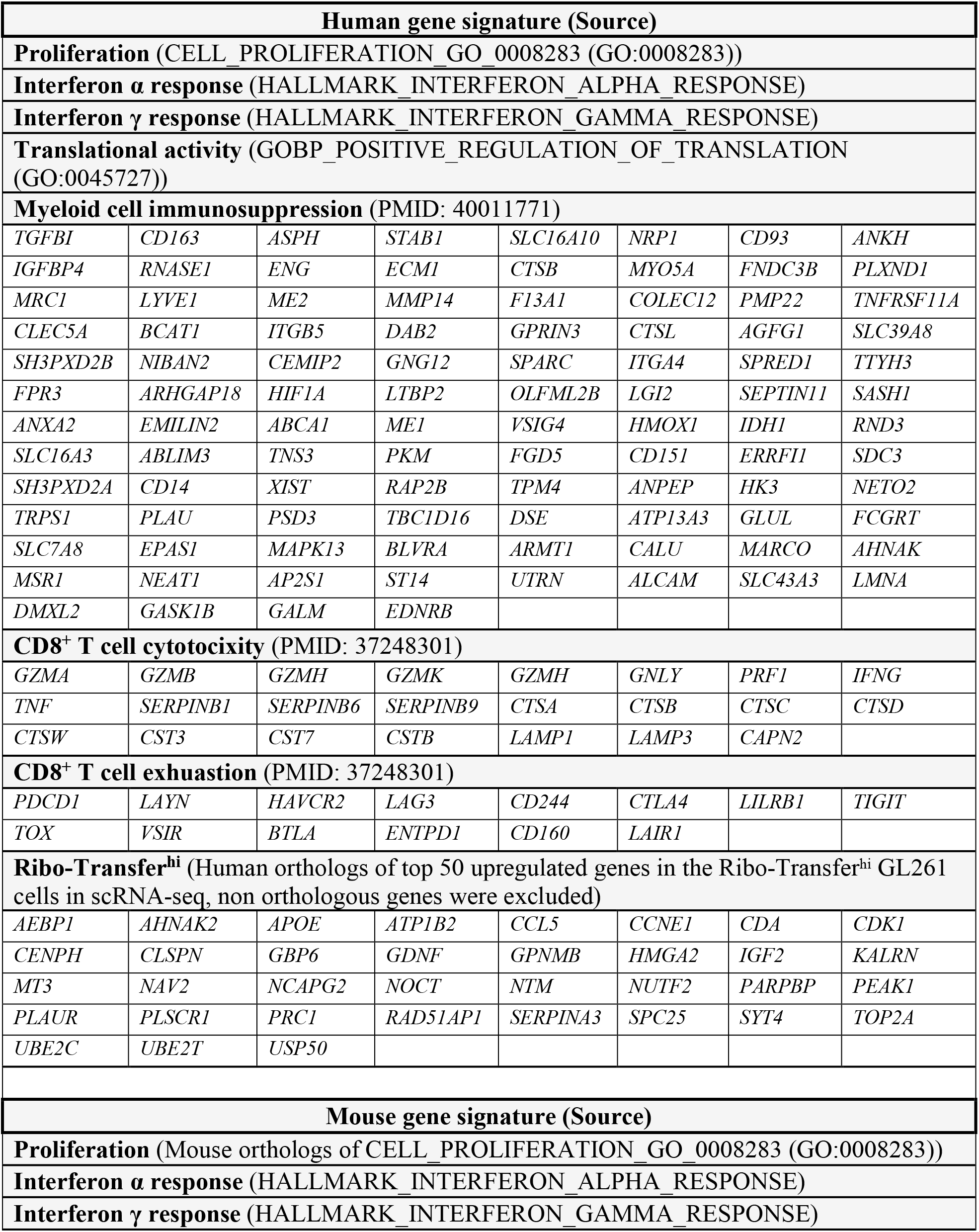
List of gene signatures.

